# The chromatin remodeling complex PRC2 safeguards cell fate in alveolar epithelial type 2 cells

**DOI:** 10.64898/2026.03.03.708777

**Authors:** Helen I. Warheit-Niemi, Jessie Huang, Kathleen C.S. Cook, Konstantinos-Dionysios Alysandratos, Sharlene Fernandes, Payel Basak, Barbara Zhao, Eitan Vilker, Carlos Villacorta-Martin, Amber Elitz, Pushpinder Bawa, Andrea Toth, Michael J. Herriges, Darrell N. Kotton, William J. Zacharias

## Abstract

Maintenance of the gas exchange surface throughout life and regeneration of the lung after injury requires tight regulation of epithelial cell fate and function. Alveolar epithelial type 2 (AT2) cells serve as the progenitors of the distal epithelium, differentiating into alveolar epithelial type 1 (AT1) cells or proliferating to maintain the quorum of AT2 cells. Here we describe the role of the chromatin regulator polycomb repressive complex 2 (PRC2) in the maintenance of AT2 cell fate in the adult alveolus. Cross-species single-cell transcriptomic analyses identified PRC2 activation in proliferative AT2 populations. PRC2 loss of function in human iPSC-derived AT2 (iAT2) cells and primary murine AT2 cells *in vitro* resulted in loss of AT2 cell state and emergence of programs reminiscent of alveolar-basal intermediate (ABI) cell states, while overexpression of the PRC2 enzymatic component EZH2 in human iAT2 cells augmented the AT2 cell program. Genetic loss of PRC2 function in the AT2 lineage in adult mice *in vivo* led to emphysematous remodeling of the lung and induced a time-dependent series of transitions of AT2 cells through an alveolar-basal intermediate (ABI) state into Krt5^+^ basal-like cells. Comparison of murine ABI cells to human disease-associated ABI cells demonstrates de-repression of canonical PRC2 targets during transition to ABI and basal-like states in human fibrosis, implicating PRC2 is a conserved regulator of AT2 cell fate. Together, these findings define PRC2 complex function during AT2 cell self-renewal as a critical guardrail for maintaining epithelial cell fate in the adult lung.

## Introduction

Lung regeneration after acute and chronic injury requires appropriate and timely activation of progenitor lineages^1^. Alveolar epithelial type 2 (AT2) cells produce and secrete surfactant lipids and proteins to maintain surface tension and are the facultative progenitors of the lung alveolus. Following injury, AT2 cells proliferate (self-renew) and differentiate into AT1 cells. While recent data has demonstrated that AT2 cells drive distal lung regeneration following injury in a process mediated by WNT and FGF signaling^2,3^, how the AT2 progenitor state is regulated at the genomic level during the injury-repair process remains poorly understood. Recent profiling studies indicate that AT2 cell progenitors are epigenetically primed for regeneration^3^ and require continuous maintenance from lineage transcription factors (TF)^4^, but the transcriptional and epigenetic factors defining this primed state are unclear. Characterizing these mechanisms will likely be necessary for development of therapeutics to recover lung alveolar function after injury.

Cell epigenomic states are defined by chromatin structure and compaction, which are mediated by methylation and acetylation of conserved histone residues, including lysine 4 and lysine 27 of histone H3 (H3K4 and H3K27). These marks regulate accessibility of chromatin to TFs, which then mediate gene expression at associated loci^5,6^. This structural regulation is conserved throughout evolution^7^ and represents a tightly controlled mechanism by which cells regulate their functional potential^8^. “Open” regions of chromatin are generally associated with transcribed genes or genes which are undergoing active repression by TFs, while “closed” regions of chromatin are less accessible. Therefore, transcription of genes in these “closed” regions requires significant change in chromatin prior to activation^9^. Throughout cell fate specification and maintenance, cells control their chromatin carefully^10^ to maintain appropriate cell fate and gene expression programs. While recent data demonstrate critical roles in the development of the respiratory epithelium for multiple epigenetic complexes including the BRG1/BRM-Associated Factors (BAF) complex^11,12^, PRDI-BF1 and RIZ homology domain containing (PRDM) 3 and 16^13^, and PRC2^14,15^, the role of epigenetic regulators in the maintenance and regeneration of the adult respiratory system, particularly in facultative AT2 progenitors, remains largely unexplored. These regulators are likely to be especially important in progenitor cells which face the daunting challenge of passing on their precise chromatin state to their progeny during self-renewing cell divisions required to maintain the progenitor pool throughout life.

Here, we show that activity of the PRC2 complex is upregulated during adult AT2 self-renewal and is required for maintenance of AT2 cell fate during adult respiratory epithelial homeostasis. The PRC2 complex is composed of the core components Suz12, Eed, and either Ezh1 or Ezh2. Suz12 functions as a structural platform for assembly of the other components, while Eed is the chromatin “reader” which identifies H3K27 residues and targets PRC2 to the correct genomic locations. Ezh1/2 are partially redundant “writer” components which catalyze the methylation of H3K27^16^. The PRC2 complex deposits and maintains H3K27me^3^ repressive marks throughout the genome^17,18^, preventing expression of genes from alternate lineages to maintain cell identity^19^. We find that genetic or pharmacological inhibition of PRC2 in human induced pluripotent stem cell-derived AT2 (iAT2) cells and primary murine AT2 progenitor organoids *in vitro* leads to loss of the AT2 cell state and emergence of markers associated with recently described aberrant states variably referred to in recent literature as transitional, intermediate, cell cycle arrested, PATS, DATP, aberrant basaloid, KRT17^+^/KRT5^−^, or alveolar-basal intermediate (ABI)^20-25^. We found that AT2-specific conditional knockout of PRC2 function *in vivo* in adult mice causes a progressive loss of AT2 cell state, with cells passing through a definable series of ABI states present in both mouse and human lung, culminating in basal cell-like identity within the alveolar epithelium. Our data demonstrate a crucial role for PRC2 in safeguarding AT2 progenitor cell fate in the adult lung, defining a targetable chromatin modifier that may be critical for both lung homeostasis and disease pathogenesis.

## Results

### Cyclical kinetics of proliferation and maturation profiles in iAT2 cells

We initially sought to identify potential regulators of human AT2 cell self-renewal using the human iPSC-derived AT2 (iAT2) cell *in vitro* model system. We have previously shown that human iAT2 cells self-renew indefinitely in defined serum-free, feeder-free AT2 culture conditions^26,27^ with serial passaging every 11-14 days. Alternatively, iAT2s can be induced to differentiate into alveolar type 1 (iAT1) cells if the culture medium is changed to AT1 promoting conditions^28^. In self-renewing AT2 conditions, we have observed an inverse association of AT2 proliferation vs. maturation genes *in vitro*^*27,29*^. To profile the dynamics of human AT2 proliferation and maturation, we first performed short-pulse EdU labeling followed by time series transcriptomic profiling by both RT-qPCR and scRNA-seq in iAT2 cells during two serial passages in AT2 cell medium (Figure 1A-B, Extended Data Figure 1A). We found the percentage of proliferating iAT2 cells (defined by EdU labeling after a 24-hour pulse) peaked between 2-5 days post-passage at 90% and decreased steadily from 7-11 days post-passage. These findings were supported by active cell cycle phase scores (S, G2/M) in transcriptomic profiling (Figures 1B-D; Extended Data Figure 1B-C). Upon the subsequent passage, the percentage of EdU^+^ iAT2 cells again increased to 90% at 2-5 days post-passage, suggesting cyclical proliferation kinetics of iAT2 cells *in vitro* (Figure 1B). Across these time points, expression of the AT2-specific transcript *SFTPC*, and AT2 maturation-associated transcripts *SFTPA1* and *SFTPA2*^30^ lagged, peaking at 7 days post-passage, following the peak of proliferation (Figure 1E-F; Extended Data Figure 1D) and corresponded to a time point when proliferation markers were decreasing (Figure 1D). These dynamics were consistent with our previously published inverse relationship between proliferation and maturation in iAT2 cells^30^. This method identified actively cycling cells for further study and indicated cyclical kinetics of proliferation and maturation that are recapitulated across serial passages in this human culture system (Figure 1F).

**Figure 1.**
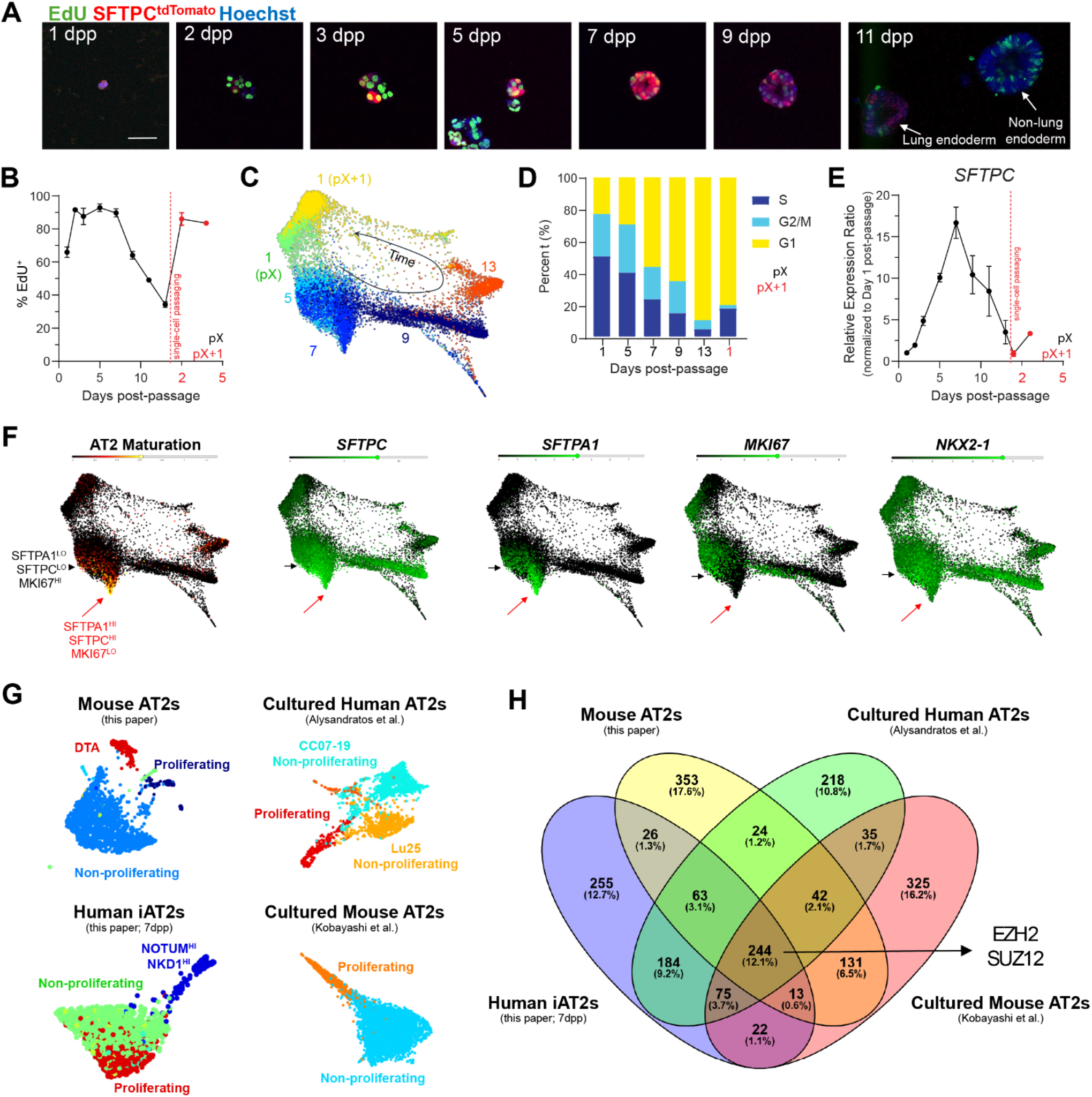
Identification of PRC2 as a potential regulator of the AT2 progenitor state. (A-B) Proliferative kinetics of iPSC-derived AT2 cells *in vitro*. (C-D) Single cell sequencing and cell cycle state of multiple stages of iAT2 growth post-passage. (E) Relative expression of SFTPC per day after passage. (F) Correlation of marker gene expression and proliferative state (black arrow) with AT2 maturation signature (red arrow) in iAT2 cells. (G-H) Evaluation of multiple new (left column) and published^24,28^ (right column) scRNAseq datasets containing AT2 cells to generate a core signature of proliferative AT2 cells (arrow, H) including PRC2 components EZH2 and SUZ12.

### Identification of PRC2 complex activation in proliferative AT2 cells

To determine transcriptomic AT2 proliferation programs shared across human and mouse species in both iAT2 cells and primary AT2 cells, we next performed a comparative analysis of 4 scRNAseq datasets: 1) our above iAT2 profiles, 2) a published profile of cultured primary human AT2 cells^27^, 3) a published profile of cultured mouse AT2 cells^23^, and 4) a new profile we prepared characterizing *in vivo* AT2 cell proliferation in mice following lineage-specific diphtheria toxin A (DTA)-induced AT2 ablation (Figure 1G; Extended Data Figure 1E). In this last model, we administered a single dose of tamoxifen to *Sftpc*^creERT2^ R26R^DTA^ animals, and identified peak proliferation based on Mki67 expression 2 days post-injury (dpi) (Extended Data Figure 1F-H). In contrast, AT2 cells in uninjured controls were largely quiescent as previously described^2,3^. We profiled the lung epithelial cells from this model at 2 dpi (Extended Data Figure 1I-M) and then compared proliferating and quiescent AT2 cells across the four datasets (Figure 1G-H). We identified 244 differentially expressed genes (DEGs) that were commonly enriched in all four models, including the expected enrichment of cell cycle-related transcripts (e.g. TOP2A, MKI67, AURKB and multiple cyclin dependent kinases (CDKs) and kinesins [KIFs]); KEGG pathway analysis showed enrichment in DNA replication and cell cycle modules (Extended Data Figure 1N). Notably, we found that multiple transcripts encoding PRC2 members, including Ezh2/EZH2 and Suz12/SUZ12 (Extended Data Figure 1O-Q) were enriched in proliferating AT2 cells across all datasets. Prediction of upstream regulators of all upregulated target genes identified Ezh2/EZH2 as the top ranked regulator (Figure 1H, Extended Data Figure 1O), further suggesting a role for PRC2 in proliferating AT2 cells.

### PRC2 is required for maintenance of the AT2 program in vitro

To functionally test the role of PRC2 in maintenance of AT2 cells, we examined effects of PRC2 loss of function in human iAT2 cells and primary murine AT2 cells *in vitro*. First, we examined the histone methyltransferase activity of EZH2, which functions in the PRC2 complex as the catalytic subunit to regulate H3K27 methylation status (Figure 2A). Using immortalized human A549 cells (Figure 2B) and iAT2 cells (Figure 2C), we found that treatment with small molecule inhibitors for EZH2 (GSK126 or 3-deazaneplanocin A [DZNep]) reduced H3K27me trimethylation at the protein level. Inhibition of EZH2 with GSK126 in iAT2 cells decreased expression of markers of AT2 differentiation and maturation, such as *SFTPC, SFTPA1*, and our previously published SFTPC^tdTomato^ reporter^31^ (Figure 2D-E). Expression of lung epithelial identity-associated transcription factor *NKX2-1* was unchanged while *SOX9* expression significantly increased following GSK126 treatment (Figure 2E). While cell count remained unchanged, proliferation-associated genes *MKI67* and *PCNA* increased following EZH2 small molecule inhibition (Figure 2F). The loss of AT2 program markers was confirmed by shRNA-mediated knockdown of *EZH2* (Figure 2G-H). In contrast, lentivirus-mediated overexpression of *EZH2* increased expression of both *SFTPC* and the SFTPC^tdTomato^ reporter in iAT2 cells (Figure 2I-J). To test the impact of EZH2 inhibition on differentiation capacity of iAT2, we generated iAT2s carrying bifluorescent NKX2-1^GFP^ and AGER^tdTomato^ reporters and exposed them to our published LATS inhibitor-containing medium that promotes iAT2 differentiation^28^. EZH2 inhibition significantly reduced iAT2 to iAT1 differentiation (Figure 2K-L).

**Figure 2.**
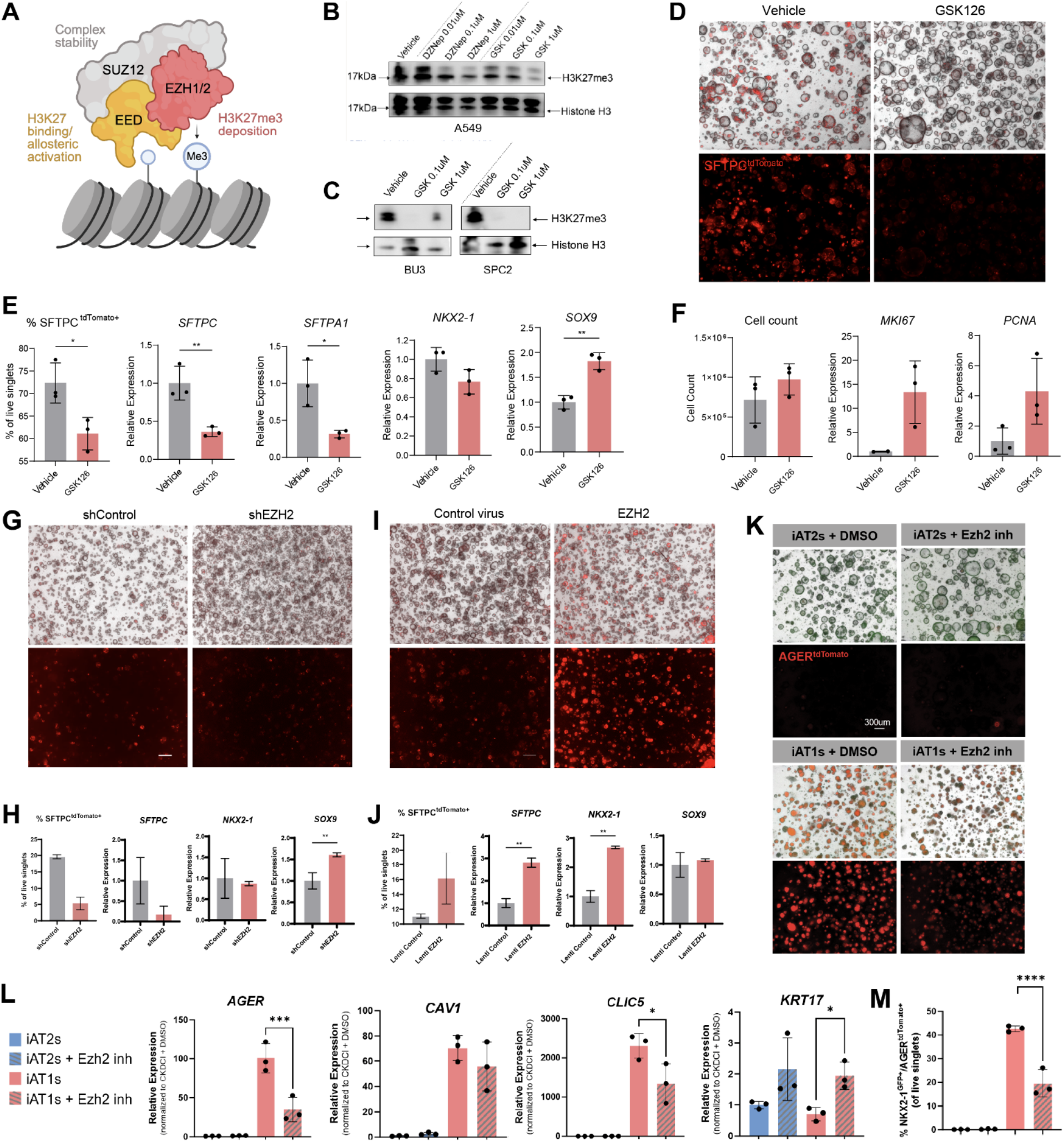
Loss of PRC2 in human AT2 progenitors results in loss of AT2 identity. (A) Schematic outlining core components of the PRC2 complex. (B-C) Confirmation that inhibition of EZH2 with GSK126 leads to reduction of H3K27me3 deposition in A549 cells (B) and two different iAT2 cell lines (C). (D) Well images showing phase-contrast and SFTPC^tdTomato^ changes with small molecule-mediated EZH2 inhibition by GSK126. (E) Loss of AT2 program in iAT2 cells after EZH2 inhibition with GSK126. Percentage of SFTPC^tdTomato^+ cells and AT2 gene expression decreases following EZH2 inhibition. (F) Evaluation of cell proliferation after GSK126 treatment. (G-J) Well images and quantification of SFTPC^tdTomato^+ cells and AT2 gene expression following shRNA-knockdown of EZH2 (G-H) or EZH2 overexpression (I-J). (K-M) Differentiation of iAT2 cells to iAT1 cells in the BU3 NGAT line show reduction in AGER^tdTomato^ reporter expression and reduced AT1 cell markers with EZH2 inhibition.

To interrogate canonical PRC2 function in primary murine alveolar organoids, we generated two independent AT2-specific conditional knockout models to delete either Ezh2 or Eed, an essential structural component required for PRC2 stability and function (Figure 2A). We generated AT2 lineage-specific tamoxifen-induced genetic ablation (conditional knockout; CKO) of PRC2 function using *Sftpc*^creERT2^ x R26R^EYFP^ or R26R^tdTom^ mice bred with Ezh2^f/f^ (Ezh2^CKO^) or Eed^f/f^ (Eed^CKO^) mice (Extended Data Figure 2A). We induced recombination with tamoxifen *in vivo* and, after 2 days, sorted Eed^CKO^ or Ezh2^CKO^ AT2 cells (Live, EPCAM^+^, EYFP^+^ cells, Extended Data Figure 2B) to establish alveolar organoids grown with supportive mesenchyme in 50% Matrigel as previously described (Extended Data Figure 2C)^3,4^. We observed reduced colony-forming efficiency in both Eed^CKO^ and Ezh2^CKO^ organoid cultures compared to controls (Extended Data Figure 2D), though both Eed^CKO^ and Ezh2^CKO^ AT2 cells generated alveolar organoids. Consistent with our human iAT2 results, both Eed^CKO^ and Ezh^CKO^ mouse organoids demonstrated loss of pro-SPC expression in AT2 cells (Extended Data Figure 2E-J) and exhibited altered morphology of RAGE^+^ AT1 cells (Extended Data Figure 2E-G). We noted increased numbers of Krt8^hi^ epithelial cells (Extended Data Figure 2H-J), consistent with acquisition of stressed or so-called “transitional alveolar cell states”^20,21,23,24,32^. Combined with our findings from human iAT2 spheroids, we concluded that cell-intrinsic PRC2 function was necessary for maintenance of the AT2 program *in vitro*.

### Loss of PRC2 function in adult AT2 cells in vivo impairs alveolar homeostasis

Based on these *in vitro* studies, we hypothesized that PRC2 activity would be functionally required to maintain the AT2 cell state during lung homeostasis *in vivo*. To test this hypothesis, we generated Eed^CKO^ and control mice in early adulthood (6-8 weeks of age) and used lineage tracing to evaluate epithelial cell identity over time during 9 months of aging (Figure 3A). *Sftpc*^creERT2^-mediated recombination was equivalent in control and Eed^CKO^ mouse lines (∼90% of total AT2 population) (Extended Data Figure 3A-D). Two weeks after tamoxifen treatment to induce Eed deletion and lineage labeling, Eed^CKO^ mice were healthy, with no significant differences in alveolar epithelial cell populations or lung structure compared to control mice (Extended Data Figure 3E-H). Evaluation at 2-, 4-, and 9-months post-tamoxifen treatment, however, showed a gradual disruption of distal lung architecture in Eed^CKO^ mice, with simplified alveolar structures, interstitial thickening, and alveolar infiltrates (Figure 3B). Nine months post-tamoxifen, mean linear intercept was significantly increased, consistent with emphysema (Figure 3B-C). We observed no difference in mortality in Eed^CKO^ mice relative to controls over the 9-month time course.

**Figure 3.**
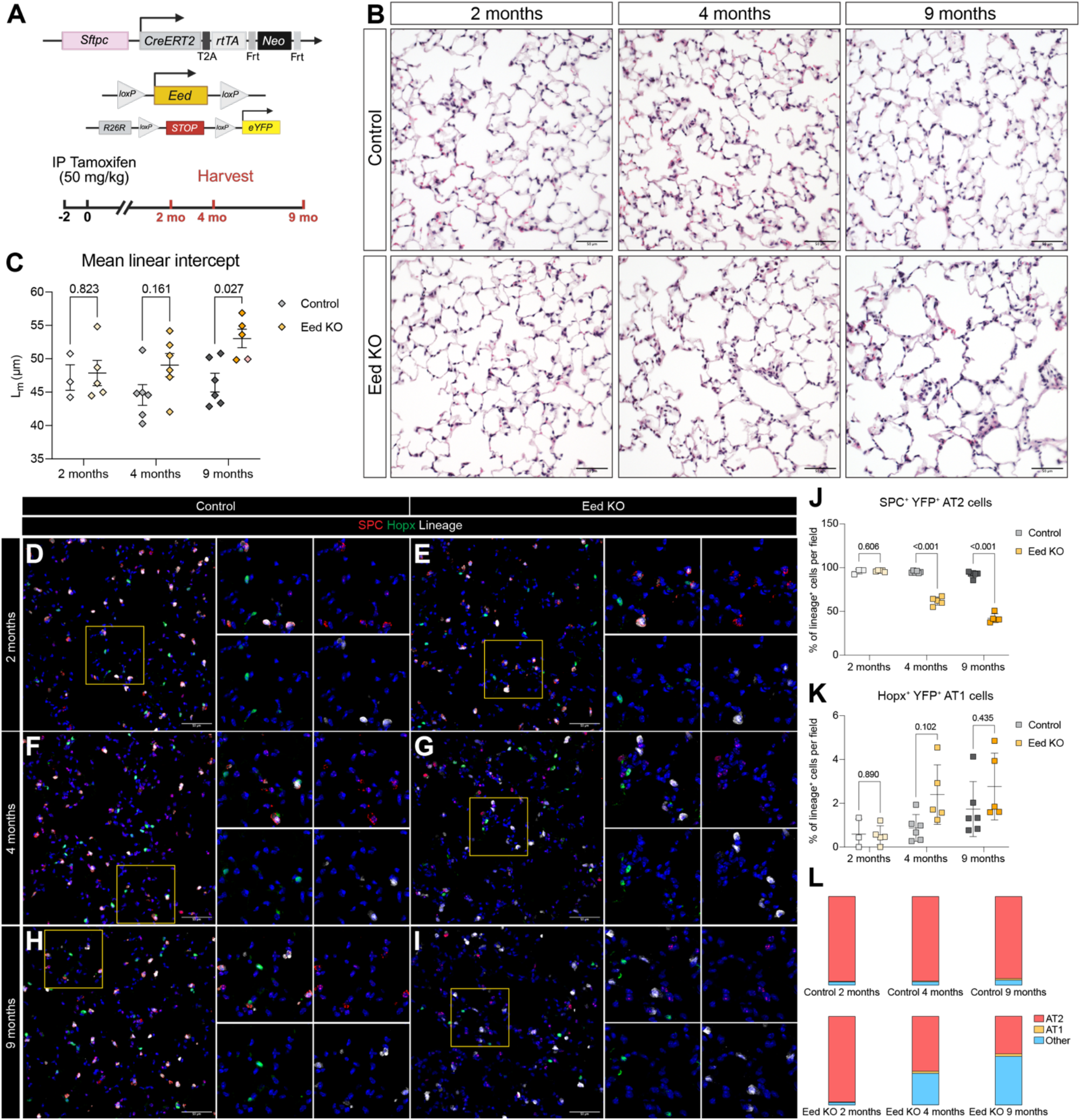
AT2 cell-specific loss of PRC2 drives spontaneous emphysema and loss of AT2 cell fate *in vivo*. (A) Schematic detailing genetics of Sftpc^creERT2^ Eed floxed mouse with fluorescent lineage reporter used for *in vivo* experiments. Mice were treated with 2 doses of 50 mg/kg tamoxifen and harvested 2-, 4-, and 9-months post-tamoxifen treatment. (B) Representative H&E images of control and Eed KO mice taken 2-, 4-, and 9-months post-tamoxifen treatment. (C) Mean linear intercept from mice described in B. Dots represent individual biological replicates from n = 3-6 mice per group. Bars represent means +/-SEM. Statistical analysis by multiple t-tests. (D-I) Representative IF images from mice described in B stained for SPC, HOPX, and YFP lineage reporter. (J) Quantification of SPC^+^ YFP^+^ AT2 cells as a proportion of total YFP^+^ cells from mice in D-I. Dots represent individual biological replicates from n = 3-6 mice per group. Bars represent means +/-SEM. Statistical analysis by multiple t-tests. (K) Quantification of HOPX^+^ YFP^+^ AT1 cells as a proportion of total YFP^+^ cells from mice in D-I. Dots represent individual biological replicates from n = 3-6 mice per group. Bars represent means +/-SEM. Statistical analysis by t-tests. (L) Stacked bar graphs representing the mean SPC^+^ YFP^+^ AT2 cells, HOPX^+^ YFP^+^ AT1 cells, and SPC^−^ HOPX^−^ YFP^+^ cells as a proportion of total YFP^+^ cells from mice in D-I.

To assess the frequencies and fates of epithelial cells within the *Sftpc*^creERT2^ lineage at each time point after Eed deletion, we performed immunostaining of mouse lung tissue sections for the AT2 marker pro-SPC, the AT1 marker HOPX, and the YFP lineage marker. Although AT2 and AT1 frequencies were unchanged at 2 months, we identified a striking time-dependent loss of lineage-labeled pro-SPC^+^ AT2 cells at 4- and 9-months post-tamoxifen (Figure 3D-J, L). AT1 cells were unchanged at all time points, likely due to the slow turnover of the distal epithelium at homeostasis^2,3^ (Figure 3D-I, K-L). These findings were not due to decreased AT2 cell proliferation, which was unchanged in Eed^CKO^ and control mice (Extended Data Figure 4A-B).

### Single cell RNA sequencing characterizes transitional cell trajectory and acquisition of basal cell-like fate following PRC2 loss of function

To better define the molecular states of cells within the Eed^CKO^ lineage over time, we generated single cell RNA sequencing (scRNAseq) libraries containing sorted YFP^+^ epithelial cells from control and Eed^CKO^ animals at 2, 4, and 9 months post-tamoxifen (Figure 4A). *The Sftpc*^*creERT2*^ WT lineage at all timepoints contained primarily AT2 cells as well as a small proportion of AT1 cells consistent with prior reports using *Sftpc*^*CreERT2*^ lines^2,3^ (Figure 4B-I). Within Eed^CKO^ mice, we identified progressive loss of AT2 cells within the *Sftpc* lineage, consistent with immunohistochemistry results (Figure 3). Instead, we identified the emergence of cells in a variety of transitional, stressed, and more proximal cell states. Within the AT2 cluster at 2 and 4 months post-tamoxifen, we detected a large fraction of cells we denoted as KO-associated AT2 cells (hereafter KO_AT2) that shared expression of AT2 markers (*Sftpc, Abca3*) with higher level expression of *Krt8*, but without significant expression of *Cldn4 or Sfn* (Figure 4B-I).

**Figure 4.**
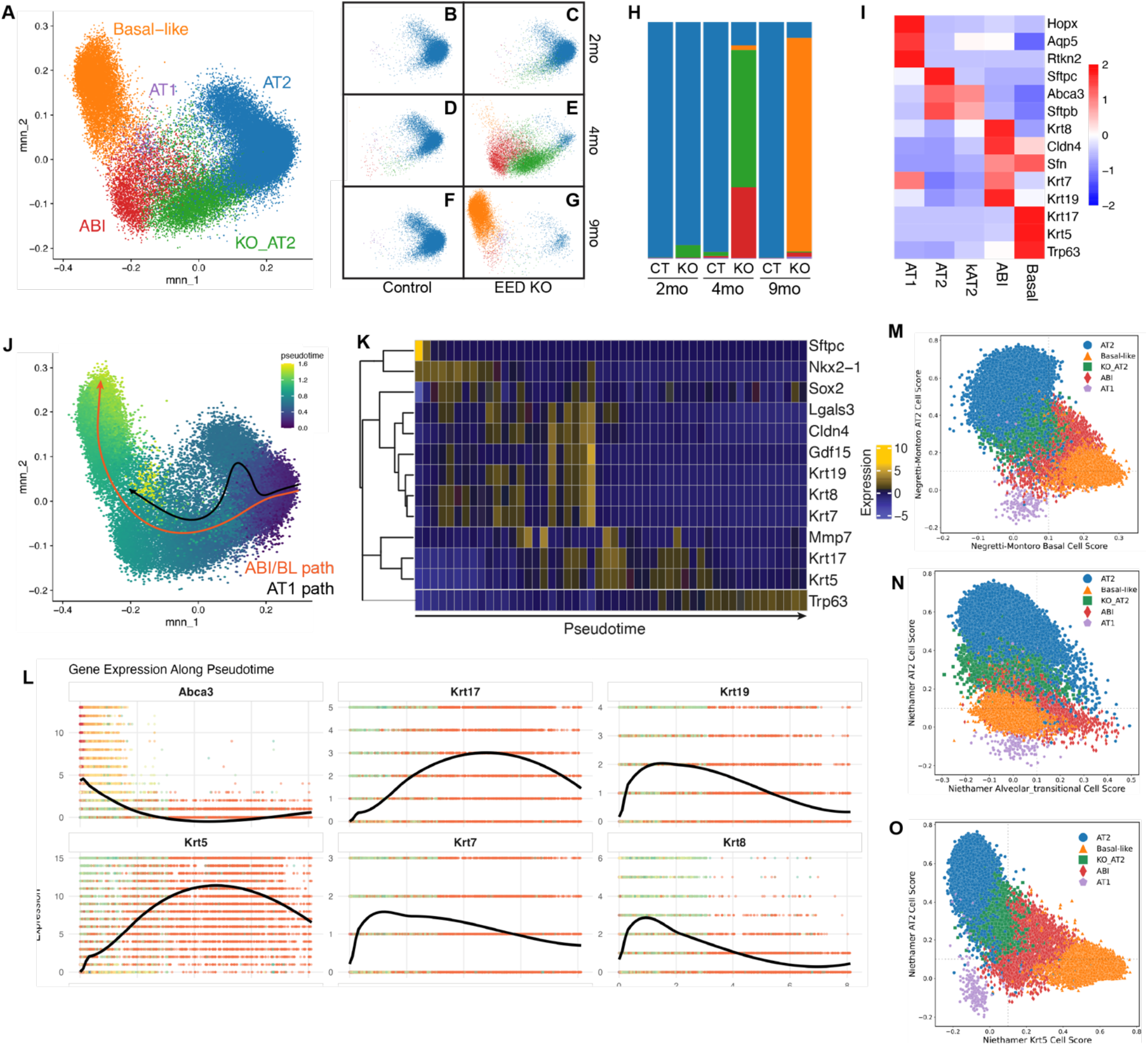
AT2 cells lacking PRC2 undergo transition to basal-like epithelia via an alveolar-basal intermediate state. A-G) UMAP projection of scRNAseq of sorted YFP+ epithelial cells from Eed^CKO^ mice at 2, 4, and 9 months. (B-G) show individual UMAP per timepoint and genotype. (H) Frequency of cell states in each timepoint. (I) Key marker gene expression in each cell state. (J) Pseudotime ordering of cell states in scRNAseq from EED^KO^ mice corresponding to temporal progression seen in A-H. (K) Heatmap of selected genes changed across the basal-like trajectory shown in (J) showing expression intensity across 50 pseudotime bins along AT2 -> basal cell differentiation. (L) Selected gene expression trajectories while undergoing alveolar to basal transition intermediates. (M-O) scTOP projections demonstrating similarity of different cell states identified in A-I to published basal cell (M), transitional cell/ABI (N) and mouse Krt5+ cell (O) states.

By 4 months post-tamoxifen, we noted emergence of cells in a state characterized by expression of multiple markers of previously described “transitional” or “alveolar-basal intermediate” cells (ABI)^22,33^. These cells have been previously defined by the combined expression of markers including *Krt8, Krt7, and Cldn4* in mice^34,35^ accompanied by diminished expression of AT2-defining markers, including *Sftpc* and *Abca3*. Human ABI cells have been further separated into KRT7^+^ ABI1 and KRT17+/KRT5-ABI2 populations^33^, with KRT17+/KRT5-cells also described as “aberrant basaloid” and associated with regions of more severe fibrosis in established fibrotic disease^22,25^. Human *in vitro* analysis has placed these intermediates on a time-dependent continuum between AT2 parents and KRT5^+^ basal cell-like descendants^33^. To date, it has remained unclear whether analogous mouse AT2-derived transitional cells are similarly competent to give rise to basal-like cells if sufficiently stressed *in vivo*. In Eed^CKO^ mice, we identified cells within the *Sftpc* lineage expressing a signature consistent with acquisition of the ABI state, including high level expression of *Cldn4, Krt7, Krt8*, and *Krt19* (Figure 4I). These cells first arose at 2 months post-tamoxifen, with a large proportion present by 4 months post-tamoxifen (Figure 4H-I). By 9 months post-tamoxifen, this population was largely absent, and instead we noted the emergence of lineage-traced basal-like cells expressing *Krt17, Krt5*, and *Trp63* (Figure 4H-I). These data are consistent with the interpretation that murine AT2-lineage cells can proceed through an alveolar-basal intermediate state to acquire basal-like characteristics in Eed^CKO^ mice. Further supporting this interpretation, pseudotime analysis of scRNAseq informed by temporal lineage tracing demonstrated a differentiation trajectory from AT2 cells to basal-like cells through KO_AT2 and ABI states (Figure 4J). This differentiation trajectory was driven by temporally ordered, state-specific gene modules which confirmed the known decrease in Nkx2-1 during AT2 to ABI transition^4^, increase in Trp53 targets including Gdf15^36^ during the KO_AT2 and ABI stages^24^, and acquisition of *Trp63* and *Krt5* late in progression from ABI to basal cells^33^ (Figure 4K-L).

To further define the cellular identity of ABI and basal-like populations observed in the Eed^CKO^ animals, we applied our published algorithm, single cell type order parameters (scTOP)^37^, to perform unbiased global transcriptomic comparisons by projecting Eed^CKO^ cell states onto reference transcriptional signatures. To that end, we used multiple independent, published reference datasets: two uninjured mouse lung scRNAseq atlases^38,39^ containing AT2, AT1, and tracheal basal cell populations, and a mouse lung injury time series dataset profiling lungs after influenza infection, which includes AT2 cells, injury-induced alveolar transitional cells, and parenchymal Krt5^+^ cells^40^. Using scTOP projection (Figure 4M-O, Extended Data Figure 5), Eed^CKO^ *Sftpc* lineage-traced basal-like cells aligned with uninjured tracheal basal cells, supporting the annotation of this cluster as “basal-like” (Figure 4M). Conversely, the Eed^CKO^ ABI population aligned with alveolar transitional cells from injured lungs (Figure 4N).

When placed on an axis comparing AT2 and Krt5^+^ reference bases (Figure 4O), Eed^CKO^ KO_AT2 and ABI cells appear as sequential intermediates between AT2 and basal/Krt5^+^ identities (Figure 4M-O, Extended Data Figure 5). Together, these analyses support the “ABI” designation and align Eed^CKO^ ABI cells with published mouse injury-induced “alveolar transitional cells”. Taken together, these scRNAseq data suggest an ordered progression of cell states from AT2 to ABI to basal-like following PRC2 loss of function.

### AT2 to basal-like cell differentiation progresses through definable transitional states in situ

To confirm these computation scoring predictions, we next performed a marker-based comparison of Eed^CKO^-derived populations to both mouse and human transitional/ABI published cell profiles via immunostaining. Prior data has shown acquisition and apparent stability of ABI states in primary human idiopathic pulmonary fibrosis (IPF) tissue, especially within heavily fibrotic alveolar regions^22,33,41^. As noted above, in human disease, these ABI cells have been referred to as ABI1 and ABI2 populations, characterized by varying levels of SFTPC (SPC), KRT8, KRT7, KRT17, and KRT5 expression, starting from a SPC^hi^/KRT8^lo^ AT2 cell state, passing through SPC^lo/^KRT8^hi^/KRT7^hi^ ABI1 and then SPC^−^ /KRT17^hi^/*KRT5*^−^ ABI2 cell states, and terminating in a KRT5^hi^/TP63^hi^/KRT17^hi^ basal cell-like state^33^. We examined whether this progression was evident in Eed^CKO^ mice by evaluating these markers 2, 4, and 9 months post-tamoxifen to quantify cells in each predicted cell state (Figure 5A-P). First, we examined the SPC and Krt8 expression in YFP^+^ lineage-traced cells and noted progressive loss of SPC^+^/Krt8^lo^ AT2 cells (Figure 5 A-C, M). The number of lineage^+^ SPC^+^/Krt8^hi^ KO_AT2 cells peaked at 2 months post-tamoxifen, steadily declining until 9 months post-tamoxifen, mirroring trends in scRNAseq data (Figure 4 B-I). The population of lineage-positive SPC^−^/Krt8^hi^ cells increased over time, illustrating differentiation away from AT2 fate.

**Figure 5.**
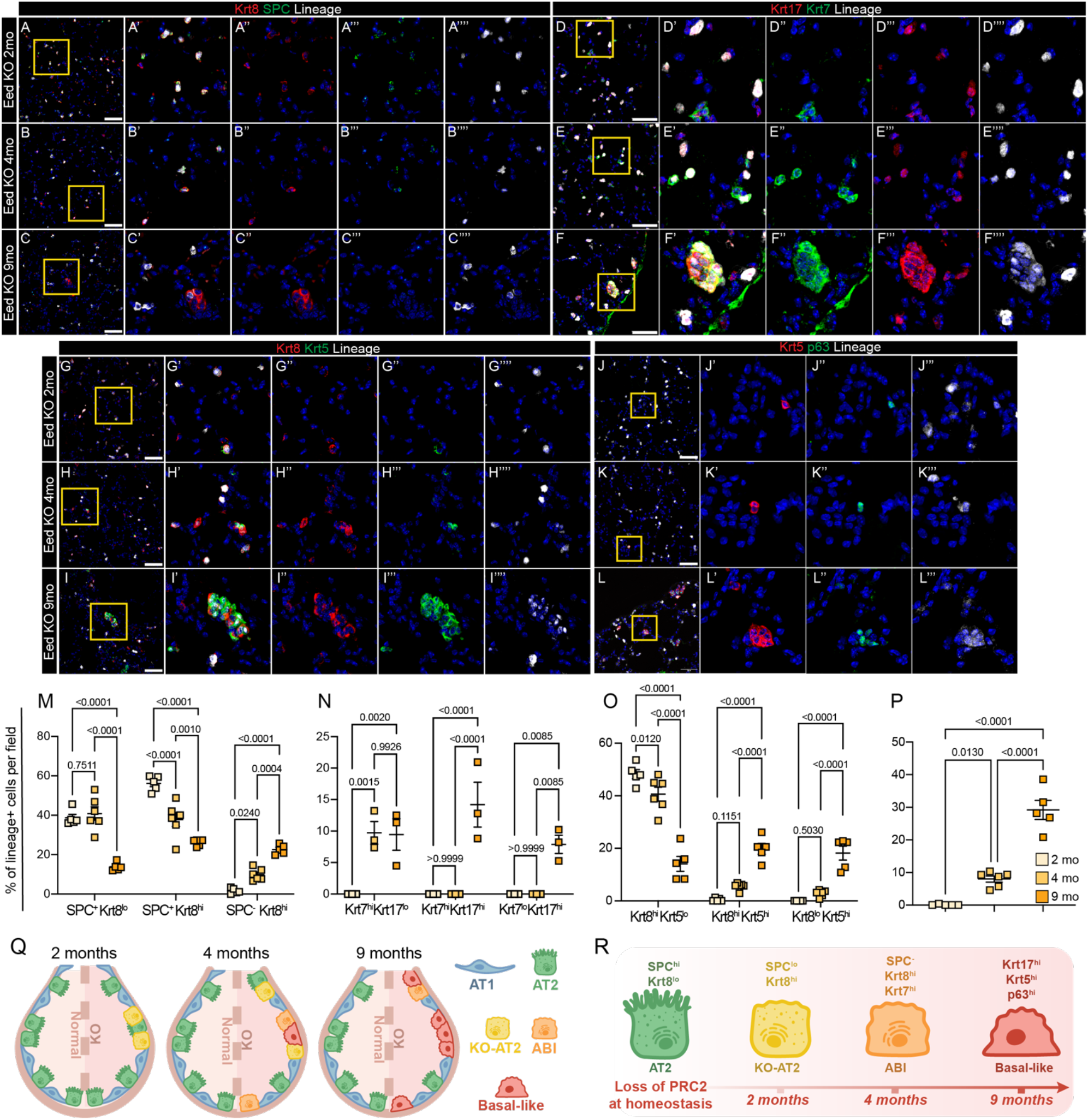
Acquisition of basal-like cell fate in murine AT2 cells lacking PRC2. (A-C) Representative IF images from Eed^CKO^ mice 2 (A), 4 (B), and 9 months (C) post-tamoxifen treatment stained for KRT8, SPC, and YFP. (D-F) Representative IF images from from Eed^CKO^ 2 (E), 4 (F), and 9 months (G) post-tamoxifen treatment stained for KRT7, KRT17, and YFP. (G-I) Representative IF images from from Eed^CKO^ 2 (E), 4 (F), and 9 months (G) post-tamoxifen treatment stained for KRT8, KRT5, and YFP. (J-L) Representative IF images from Eed^CKO^ mice 2 (I), 4 (J), and 9 months (K) post-tamoxifen treatment stained for KRT5, P63, and YFP. (M-O) Quantification of stainings shown in A-C (M), D-F (N), and G-I (O). (P) Quantification of KRT5^+^ P63^+^ YFP^+^ alveolar cells as a proportion of total YFP^+^ cells from mice in J-L. (Q-R) Model representing temporal accumulation and markers (both RNA and protein) of cell states identified in Figures 4 and 5. Dots represent individual biological replicates from n = 5-6 mice per group. Bars represent means +/-SEM. Statistical analysis by one-way ANOVA with pre-specified multiple comparisons. Scale bars = 50uM.

Evaluation of the ABI-associated marker KRT7 (Figure 5D-F, N) showed initial appearance of KRT7^hi^ cells at 4 months post-tamoxifen, consistent with transition to an ABI cell state. Protein expression of the more basal-like marker KRT17 was undetectable in the distal lung until 9 months post-tamoxifen, when we detected KRT7^hi^/KRT17^hi^ and KRT7^lo^/KRT17^hi^ cells, consistent with transition from ABI to basal-like cells (Figure 5D-F, N). We measured the transition from ABI to lineage^+^ basal-like cells by co-staining for KRT8 and KRT5 in Eed^CKO^ mice. The number of lineage^+^ KRT8^hi^/KRT5^lo^ ABI cells declined over time while the numbers of both KRT8^hi^/KRT5^hi^ and KRT8^lo^/KRT5^hi^ lineage^+^ cells increased (Figure 5 G-I, O, Extended Data Figure 6), confirming the temporal transition from ABI to basal-like with a second set of protein markers. Organoids derived from Eed^CKO^ AT2 cells exhibited increased expression of KRT8 and KRT5 after 28 days of growth in culture, suggesting that recurrent cell proliferation could promote more rapid transition of cells through this series of intermediates (Extended Data Figure 6). These patterns of cytokeratin expression mirror the results of the scRNAseq, both in timing of cell state transition (Figure 4H) and in dynamics of gene expression of Krt8, Krt7, Krt17, and Krt5 (Figure 4L). As confirmation of basal-like identity *in situ*, we evaluated expression of the basal cell transcription factor P63, identifying lineage^+^ Eed^CKO^ cells co-expressing KRT5 and P63 (Figure 5 I-L). Together, these data suggest that loss of PRC2 in AT2 cells causes differentiation to a basal cell fate within the alveolus through temporally and molecularly definable intermediate states (Figure 5Q-R).

### Loss of function in PRC2 is a common mechanism underlying alveolar to basal transition inmouse and human

To evaluate the similarity of the observed transition from AT2 to basal cells in mouse and human, we reanalyzed the published dataset^33^ demonstrating human AT2 to basal cell transition in primary 3D organoid co-culture with supportive mesenchyme. We compared the trajectory of marker gene expression for our panel of cytokeratin markers, as well as expression of canonical PRC2-inhibited target genes during AT2 to basal differentiation (Figure 6A-D). Notably, both mouse and human AT2 cells passed through similar, definable molecular states at single cell resolution, with upregulation of similar gene programs during progression to basal-like cells. Analysis of expression of canonical PRC2-silenced target genes, some of which are senescence-associated and are upregulated in disease contexts in humans, demonstrated similar upregulation in both mouse and human data, including increased expression of *Cdkn1a/CDKN1A* (p21) and Cdkn2a/CDKN2A (p16). (Figure 6B,D).

**Figure 6.**
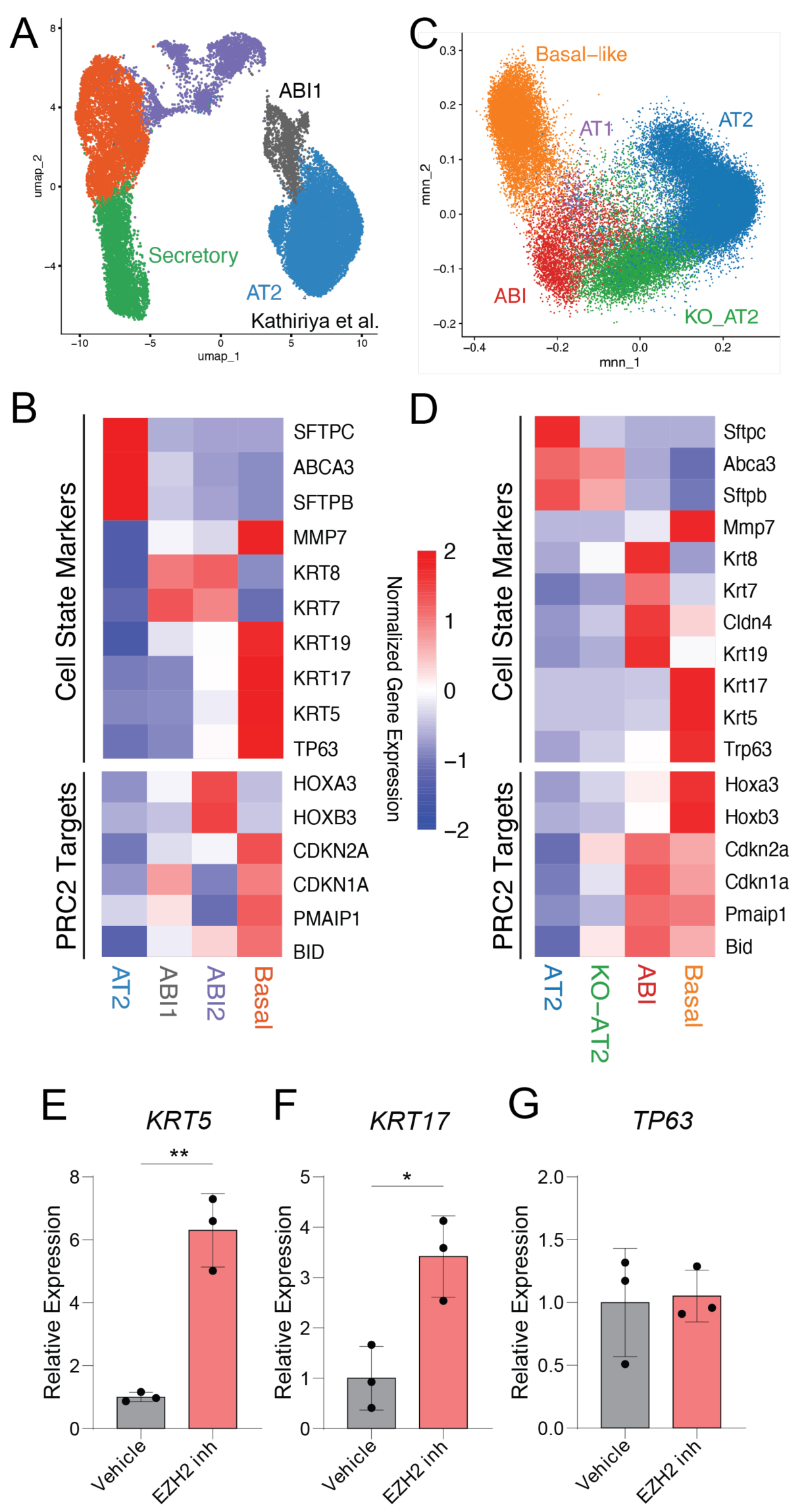
Murine and human AT2 cells transition through analogous ABI states under the control of PRC2. (A-B) Re-analysis of data from Kathiriya et al^33^, showing cell populations in transdifferentiation of human primary AT2 cells to basal and secretory identities *in vitro*. UMAP (A), cell state marker genes (per original manuscript and this study), and expression of canonical PRC2 repressed targets (increased expression with decreased PRC2 function) (B). (C-D) Mouse Eed^CKO^ and control scRNAseq data, with display mirroring human including UMAP (reproduced from Figure 4) (C), cell state marker genes and PRC2 target gene expression (D). (E-G) QPCR for markers of ABI and basal cells in iAT2 after treatment with EZH2 inhibitor GSK126. Statistical significance determined by t-test, *=p<0.05, ** p<0.01, each dot showing a single replicate; expression level normalized to vehicle.

Consistent with this model, inhibition of EZH2 in serum-free, feeder-free culture of pure human iAT2s resulted in increased expression of *KRT5* and *KRT17* (Figure 6E-F) and decreased expression of SFTPC (Figure 2E), emphasizing the importance of PRC2 in preventing AT2 to basal cell differentiation in human cells (Figure 6E-G). These data correspond to computational models of human disease which previously implicated EZH2 function in the pathogenesis of IPF^41^. Taken together, these data demonstrate a conserved mechanism whereby PRC2 activity constrains and safeguards AT2 cell fate during self-renewal. Further, these data support the concept that both mouse and human AT2 cells can differentiate into basal-like cells via a conserved mechanism dependent on control of the cellular epigenetic state.

## Discussion

We show here that the loss or inhibition of PRC2 subunits leads to loss of the AT2 program in human and murine AT2 cells, with consequent gain of a defined series of cytokeratin-expressing intermediates that culminates in a basal cell-like state. We speculate that the enrichment of PRC2 function in cycling AT2 cells identified in our study is due to the need for AT2 progenitors to safeguard or “remember” their alveolar identity during cell division, and the progressive fate drift seen *in vivo* in Eed^CKO^ mice represents time-dependent, recurrent loss of fate across several homeostatic cell divisions. These data demonstrate that AT2 cell fate is actively maintained throughout adult life, with specific cell-intrinsic and cell-extrinsic signals closely restricting the identity, self-renewal, and differentiation of AT2 cells. This evolutionarily conserved regulation of AT2 cell identity is disrupted in multiple human diseases, with emergence of disease-associated cell states on the path from AT2 to basal cells. Our findings suggest that basal cells may be a “default” cell identity for lung epithelial cells and imply that chronic injury during the pathogenesis of lung disease causes chronic epigenetic stress. If true, the epithelial lesions found in diseased tissue may represent accumulation of cell stress which causes loss of AT2 cell fate, contributing to clinical manifestations of chronic lung disease. Our model is supported by data showing disruption of PRC2 target gene expression including CDKN1A and CDKN2A in diseased epithelial states in humans^25,33,35,41^.

Our data add to emerging literature demonstrating a key role for active chromatin regulation in lung epithelium in development and throughout adulthood. PRC2 has been previously implicated in lung epithelial differentiation, wherein loss of function of Ezh2 in early endodermal progenitors led to ectopic and premature appearance of basal cells in the developing airway epithelium but not in alveolar regions^15^. In more recent work, Ezh2 loss of function in developing alveolar cells reduced AT2 to AT1 differentiation^42^. Ezh2 is a predicted target of the microRNA *let7*, recently shown to maintain distal lung epithelial fate and reduce fibrotic risk following bleomycin, orthogonally implicating Ezh2 regulation in alveolar cell fate specification^43^. Recent data have demonstrated a key role for epigenetic mechanisms underlying AT2 vs. AT1 differentiation^44^, where PRDM3/16 function is required for AT2 specification^13^ and canonical BAF complex activity is required for AT1 specification^11^. Both of these chromatin-associated complexes interact with Nkx2-1, which interacts with distinct loci in AT2 vs. AT1 cells^45^ and are required to maintain the AT2 progenitor state during lung homeostasis^4^. Future work will be needed to investigate how cell-specific TF interactions with epigenetic regulators underlie other aspects of cell fate transitions along the airway-alveolar axis.

The present data support a model of alveolar epithelial cell state control with significant therapeutic implications. Given that AT2 cells can transition to basal cells, and extensive evidence suggests similar plasticity between alveolar and secretory cells in multiple injury models^46-49^, the epithelial axis of differentiation appears to be highly plastic. Extant data suggests that distal cells are more capable of entering PATS- and ABI-like states than proximal cells, which implies that distal differentiation may require more coordinated gene expression control, or that disrupting established AT2 state via loss of genome-wide regulators such as PRC2 is more feasible than activation of specific alveolar cell differentiation mechanisms.

Finally, our study identified clear intermediate cell states between AT2 and basal cell fates in mice, which are abundant in human disease, suggesting a potential for therapeutic targeting. A next step would be identifying specific, conserved targets defining each disease-associated intermediate state. One key future direction suggested by our data is to define PRC2 partners in each intermediate cell state and identify epigenetic factors that promote distal fate acquisition to balance PRC2 activity. Given that many drugs targeting epigenetic complexes exist in therapeutic pipelines, focusing on chromatin regulation during lung health and disease presents an opportunity for development of pro-regenerative therapies in lung disease.

## Acknowledgements

The authors would like to thank the Single Cell Genomics Facility (especially Kelly Rangel and Shawn Smith), Bio-Imaging and Analysis Facility (especially director Matt Kofron), Genomics Sequencing Facility, and the Research Flow Cytometry Facility of the Cincinnati Children’s Research Foundation; the Harvard University Single Cell Core (especially Mandovi Chatterjee), the Boston University Single Cell Sequencing Core (especially Yuriy Aleksyeyev), and the Boston University Flow Cytometry Core (especially Brian Tilton), all of whom have provided extensive technical support.

## Author Contributions

Conceptualization – HWN, JH, DNK, WJZ

Data acquisition – HWN, JH, KC, KDA, SF, PB, BZ, CVM, AE, PB, AT, MH

Data analysis – HWN, JH, KC, KDA, SF, PB, BZ, CVM, AE, PB, AT, MH, DNK, WJZ

Supervision – DNK, WJZ

Writing – original draft – HWN, JH, KC, DNK, WJZ

Writing – review and editing - all authors

## Materials and Methods

### Human induced pluripotent stem cell (iPSC)-derived AT2 cells

All human iPSC work was conducted under regulatory approval of Boston University’s Institutional Review Board. The generation of iPSC-derived AT2 (iAT2) cells by directed differentiation was performed on published human iPSC lines (SPC2-ST-B2, BU3 NGST; available from www.kottonlab.com), which have either a NKX2-1^GFP^ and/or a SFTPC^tdTomato^ reporter to allow for monitoring of the AT2 program in culture, as previously described^26,30,50^. Briefly, lung progenitors were sorted (MoFlo Astrios EQ) at day 15 of differentiation and resuspended in growth factor-reduced Matrigel (Corning) at a density of 400 cells/μL Matrigel in CK+DCI medium, consisting of complete serum-free differentiation medium base supplemented with 3 μM CHIR99021, 10 ng/mL recombinant human KGF, 50 nM dexamethasone (Sigma), 0.1 mM 8-Bromoadenosine 3′,5′-cyclic monophosphate sodium salt (Sigma), and 0.1 mM IBMX (Sigma). 10 μM Y-27632 was added to CK+DCI for 48-72h after each passage. The resulting spheres were single-cell passaged and replated into Matrigel on Day 30 of differentiation, followed by a brief 3-to 4-day CHIR99021 withdrawal to achieve iAT2 maturation. Cells were subsequently sorted on day 45 of differentiation on SFTPC^tdTomato^+, and maintained in CK+DCI through serial passaging every 10-14 days. Purity was monitored at every passage by flow cytometry for SFTPC^tdTomato^ expression.

### EdU labeling of iAT2 cells in vitro

iAT2 cells were treated with EdU for 24h, as optimized by time course labeling, prior to each time point collection using the Click-iT™ EdU Cell Proliferation Kit (ThermoFisher) per manufacturer’s instructions. EdU-labeled cells were processed for flow cytometry or whole-mount immunostaining. For flow cytometry, iAT2 cells were dissociated using 2mg/mL Dispase II, then 0.05% trypsin-EDTA, and fixed and processed per manufacturer’s instructions. EdU-labeled cells were flowed (Stratedigm) and assessed for EdU percentage. For whole-mount immunostaining, spheroids were released from Matrigel using 2mg/mL Dispase II, and fixed, blocked, and stained as previously described^51^. Stained spheroids were mounted on cavity slides, and imaged on a confocal microscope (Zeiss LSM 710).

### EZH2 modulation in human A549 cells and iAT2 cells

For EZH2 inhibition experiments, A549 cells and iAT2 cells were treated with 0.01-1µM GSK126 or DZNep for 7d. For Western blotting, cells were dissociated according to their respective protocols (A549 cells with 0.05% trypsin-EDTA, iAT2 cells with Dispase II and trypsin-EDTA), and lysed using RIPA buffer (50 mM Tris-HCl pH 7.4, 150 mM NaCl, 1% NP-40, 0.5% sodium deoxycholate, 0.1% SDS) supplemented with protease inhibitor, incubated on ice for 15 minutes, and clarified by centrifugation at 15,000g for 20 minutes at 4°C. The supernatant containing total protein was collected, and protein concentration was determined using a BCA assay (Pierce) according to the manufacturer’s instructions. Equal amounts of protein were denatured in Laemmli sample buffer, boiled at 95°C for 5 minutes, and subjected to SDS-PAGE followed by immunoblotting for H3K27me3 (1:1000; Cell Signaling #9733) and total Histone H3 (1:1000; Cell Signaling #9715).

### Ethics/Animals

All animal use was done in accordance with CCHMC and BU IACUC protocols. The following mouse strains were maintained on a mixed CD-1 – C57BL/6 background and used for all murine *in vivo* and *in vitro* experiments: *Sftpc*^creERT2^ R26R^eYFP^ *Eed*^fl/fl^ (Eed^CKO^), *Sftpc*^creERT2^ R26R^tdT^ *Ezh2*^fl/fl^ (Ezh2^CKO^), *Sftpc*^creERT2^ R26R^eYFP^ (control). *Sftpc*^creERT2^ R26R^DTA^ was maintained on a C57BL/6 background. To induce Cre recombination, male and female 8-12 week old mice were given two intraperitoneal (IP) doses of 50 mg/kg tamoxifen (Sigma, T5648). *Sftpc*^creERT2^ R26R^DTA^ mice and littermate controls were given one single IP dose of 100 mg/kg tamoxifen. Prior to use, tamoxifen was dissolved in 100% ethanol and diluted in corn oil (Sigma, C8267) at a 1:9 ratio for a final stock concentration of 10 mg/mL. At specified experimental timepoints, mice were anesthetized with an IP dose of ketamine/xylazine and euthanized via cervical dislocation before the removal of the lungs.

### Histology

Following euthanasia, the chest cavity was exposed, and the right ventricle of the heart was perfused with 10 mL of cold 1X phosphate buffered saline (PBS) (Gibco 10010023). A 22G Exel Safelet catheter (Fisher, 14-841-20) was inserted into the trachea and approximately 1-2 mL of 2% paraformaldehyde (PFA) was instilled into the lungs until all lobes were inflated. The catheter was removed, trachea tied off with surgical suture, and lungs removed whole and fixed in 2% PFA for 24 hours. After fixation, lung lobes were dissected out and tissue was dehydrated through an ethanol gradient, embedded in paraffin, and cut into 5 um sections. Hematoxylin and eosin staining was performed to evaluate lung morphology. Following sodium citrate (10 mM, pH 6.0) antigen retrieval, immunohistochemistry and immunofluorescence was performed using the primary antibodies described in Supplemental Table 1. Tissue sections were incubated with primary antibodies overnight at 4 degrees C followed by either a two-hour incubation at room temperature with fluorescently conjugated secondary antibodies or amplification using the ImmPRESS HRP Horse anti-Rabbit IgG Polymer Detection Kit (Vector Labs (MP-7401-50) and subsequent application of TSA fluorophores (Akoya Biosciences) (described in Supplemental Table 1). Autofluorescence was quenched using the Vector TrueVIEW Autofluorescence Quenching Kit (Vector, SP-8500-15), samples were stained with DAPI (Invitrogen, D1306), and coverslips were mounted onto slides with Prolong Gold antifade mounting medium (Invitrogen, P36930).

### Imaging and quantification

Brightfield images were taken using a Nikon upright widefield microscope with 20X objective. Immunofluorescent images were taken on a Nikon A1 inverted LUNV with 20X and 40X water objectives. Five fields per animal were chosen randomly for image analysis. Cell types of interest were manually counted, quantified as a proportion of total lineage^+^ cells per field, and averaged per animal. Mean linear intercept was quantified as previously described^30^.

### Mouse lung digestion and single cell suspension

Single cell suspensions from murine lungs were generated as previously described^6^. Following euthanasia, the chest cavity was exposed, and the right ventricle of the heart was perfused with 10 mL of cold 1X PBS. Gross airways and non-pulmonary tissue were dissected *in situ* and lung lobes were removed and submerged in ice-cold PBS. Lung tissue was minced with scissors in an empty gentleMACS C Tube (Miltenyi 130-096-334) before adding 5 mL of 37°C digest buffer (dispase (Corning 354235), DNase1 (GoldBio D-301), collagenase type 1 (Gibco 17100017), and 1X PBS). The C tubes were placed on a gentleMACS Octo Dissociator with Heaters (Miltenyi Biotec, 130-096-427) with the protocols “m_lung_01_02” (36 s) run twice, “37C_m_LIDK_1” (36 min 12 s) run once, and “m_lung_01_02” (36 s) run once again. The samples were then passed through a 70 uM filter, centrifuged at 800g for 8 minutes at 4°C, aspirated, and incubated in room temperature RBC Lysis Buffer (eBioScience 00-4333-57) for 5 minutes. Supernatant was aspirated following centrifugation at 500g for 5 minutes at 4°C, and the cell pellet was resuspended in cold MACS Buffer [AutoMACS Rinsing Buffer (Miltenyi 130-091-222) + MACS BSA Stock Solution (Miltenyi 130-091-376)] before being passed through a 40 uM filter and washed with 5 mL of MACS Buffer. Cells were centrifuged at 500g for 5 minutes at 4°C and resuspended in a final volume of 5 mL to generate a single cell suspension.

### FACS preparation and processing

Cells were centrifuged, aspirated, and incubated in Fc Receptor Binding Inhibitor Polyclonal Antibody (eBioScience 14-9161-73) diluted 1:100 in MACS Buffer for 10 minutes at room temperature. Cells were centrifuged, supernatant was removed, and cells were incubated in CD326 APC antibody (Invitrogen 17-5791-82) diluted 1:100 in MACS Buffer at room temperature for 10 minutes in the dark. Following centrifugation and aspiration, cells were washed with 5 mL of cold MACS Buffer, then centrifuged again. Supernatant was removed and cells were incubated with Fixable Viability Dye eFluor™ 780 (eBioScience 65-0865-14) diluted 1:1000 in MACS Buffer for 15 minutes at room temperature in the dark. Cells were then centrifuged and washed 2-3X with 5 mL of cold MACS Buffer before being resuspended in a cell count adjusted volume of MACS Buffer and strained through 35 uM filter lids of polystyrene FACS tubes on ice (Corning 352235) for sorting.

Using single-stain compensation beads (Invitrogen 01111142), gating was adjusted to remove debris and doublets using a BD FACSAria Fusion or BD FACSymphony cell sorter fitted with a 100 uM nozzle. The live/CD326^+^/eYFP^+^ (AT2) population was sorted into a 1.5 mL tube pre-coated in 500 μL of cold SAGM (Lonza CC-3118) organoid media (see “Organoid media” section below) to preserve cell viability. For this protocol, the yield was approximately 2×10^5^ AT2 cells per mouse.

### Fibroblast stock preparation

Murine fibroblast stocks were generated following previously reported methods^6^. Briefly, 4-week C57BL/6J mouse lungs were harvested, digested, and a single cell suspension was achieved using techniques described above. Cells were centrifuged at 500g for 5 minutes at 4°C, supernatant was aspirated, and cells were washed three times with cold MACS Buffer. The cell pellet was resuspended in 10 mL of fibroblast media (DMEM/F-12 [Gibco, 11320-033], Antibiotic-Antimycotic [Gibco 15240-062, final concentration 1x], and Heat Inactivated Fetal Bovine Serum [Corning 35-011-CV, final concentration 10%]), distributed on a 10 cm round tissue culture plate (approximately 1 mouse lung per plate), and non-adherent cells were removed through a media change 2–4 hours post-plating.

Cells were passaged at 80% confluency to P3 by removing the media from each plate, washing the cells with 5 mL of DPBS, trypsinizing them with 3 mL of 0.25% Trypsin-EDTA (Gibco, 25200-056) at 37°C for 7 minutes, adding 5 mL of fibroblast media to each plate to halt enzymatic activity, manually pipetting and using a cell scraper to further dissociate the cells, and transferring them to a 15 mL conical tube. Cells were then centrifuged at 500g for 5 minutes at 4°C, the supernatant was aspirated, and the cell pellet was resuspended. For a 1:3 split, cells were resuspended in 6 mL fibroblast medium and 2 mL were transferred to plates containing 6 mL of fibroblast media. Once 80% confluency was reached at P3, cells were washed, trypsinized, and centrifuged as above, and resuspended in 1 mL of freezing medium (90% FBS, 10% DMSO) before being transferred into a cryovial (one plate/cryovial) and housed within a Mr. Frosty Cryogenic Freezing Container (Nalgene, 5100-0001) filled with isopropyl alcohol. The Mr. Frosty was placed in a −80°C freezer overnight and samples were moved to liquid nitrogen for long-term storage.

For the use of frozen fibroblast stocks in organoids, 48-72 hours prior to use in organoids, cells were rapidly thawed from liquid nitrogen and resuspended in a 10 mL of warm fibroblast media in a 15 mL conical tube. Cells were centrifuged at 500g for 5 min at 4°C, supernatant was aspirated, and cell pellet was resuspended in 2 mL of fibroblast media. Cells were plated on a 10 cm tissue culture plate containing 6 mL of fibroblast media, and media was changed after 24-48 hours to encourage adherence and expansion of cells.

### Mouse organoid plating

After sorting, AT2 cells were centrifuged at 500g for 5 minutes at 4°C, counted with 0.4% Trypan blue solution (Gibco 15250061), and resuspended in SAGM. Previously seeded and cultured fibroblasts (see above “Fibroblast generation and seeding”) were washed with 5 mL of DPBS (Gibco 14190144), trypsinized with 3 mL of 0.25% trypsin-EDTA (Gibco 25200056) and allowed to incubate at 37°C for 7 minutes. To neutralize the trypsin, 5 mL of DMEM/F12 media (Gibco 11320-033 supplemented with 10% FBS [Corning MT35011CV] and Antibiotic-Antimycotic [Gibco 15240062]) was added. Dissociated fibroblasts were centrifuged at 500g for 5 minutes at 4°C, counted with 0.4% Trypan blue solution, and resuspended in SAGM. Per well, 5,000 AT2 cells and 50,000 fibroblasts were combined in a volume of 45 μL SAGM per well. Ice-cold Matrigel GFR Membrane Matrix (Corning 356231) was added to the epithelial and fibroblast mixture at a 1:1 ratio and carefully mixed to prevent bubble formation while on ice. 24-well companion plates (Corning 353504) and the desired number of transwells (Corning 353095) were assembled and kept on ice throughout the plating process. To plate, 90 μL of the final cell and Matrigel mixture was carefully added to the center of each chilled transwell in the companion plate before being incubated at 37°C for 10-15 minutes to allow the Matrigel mixture to set.

Finally, 500 μL of SAGM supplemented with ROCK Inhibitor/Y-27632 dihydrochloride (Sigma Y0503) at a final concentration of 10 uM was fed beneath each transwell. ROCK inhibitor-supplemented SAGM media was added only to the initial 48 hours of culture and all ensuing media changes were completed with regular SAGM media. Organoids were cultured at 37°C with 5% CO_2_ for 28 days with media changes every 48 hours.

### Mouse organoid growth media

To generate SAGM media for murine lung alveolar organoids, SABM Small Airway Epithelial Cell Growth Basal Medium (Lonza, CC-3119) was spiked with the following additives: SAGM Small Airway Epithelial Cell Growth Medium SingleQuots Supplements and Growth Factors (Lonza CC-4124, using only the BPE [2 mL], Insulin [0.5 mL], Retinoic Acid [0.5 mL], Transferrin [0.5 mL], and hEGF [0.5 mL] aliquots), Heat Inactivated Fetal Bovine Serum (Corning, 35-011-CV, final concentration 5%), Antibiotic-Antimycotic (Gibco, 15240-062, final concentration 1x), and Cholera Toxin from *Vibrio cholerae* (Sigma, C8052, final concentration 25 ng/mL). Media was aliquoted into 50 mL quantities and kept at 4°C until use.

### Mouse organoid retrieval and whole-mount immunostaining

After 28 days of culture, mouse organoids were liberated from the 3D matrix for wholemount immunostaining. For all steps in the isolation procedure prior to fixation, plasticware (pipette tips, plates, conical tubes) was pre-coated with 1% bovine serum albumin (Sigma A3294) solution in PBS to prevent organoids from sticking. Culture media was removed from the plate wells, and they were briefly washed with 1X PBS. To isolate the organoids, the plate was kept on ice and 500 μL of ice-cold cell recovery solution (Corning 08-774-405) was added to the inside of each Transwell. The Matrigel was mechanically disrupted with a pre-coated wide bore P1000 tip (Midsci 4396-SF) before being transferred to a standard 24-well tissue culture plate (Fisher FB012929) on ice. The process was repeated with an additional 250 μL of ice-cold cell recovery solution to recover as many organoids from the 3D matrix as possible. The plate was then incubated on ice on a vigorously shaking orbital shaker (100 rpm) for 60 minutes to dissolve the 3D droplets. The contents of each well were gently resuspended with a pre-coated wide bore tip and transferred to a pre-coated 15 mL conical tube. The wells were washed with additional 1% PBS-BSA to recover all organoids. Wells of organoids with the same condition were pooled together. The conical tubes were centrifuged at 70g for 5 minutes at 4°C, supernatant was removed, and the pellet was resuspended in 1 mL of 4% paraformaldehyde and incubated at 4°C for 45 minutes. If proceeding directly to blocking and immunolabeling, organoids were permeabilized by adding 9 mL of cold 0.1% Tween-20 (Sigma P1379) in PBS and incubated overnight at 4°C. If not proceeding immediately to the blocking and immunolabeling step, the organoids were centrifuged at 70g for 5 minutes at 4°C, supernatant was removed, and the pellet was resuspended and stored in organoid washing buffer, a solution of 0.2% BSA and 0.1% Triton X-100 in PBS. In our experience, fixed organoids can remain suspended in organoid washing buffer for up to 1 year with no observed effect. To block, organoids were centrifuged at 70g for 5 minutes at 4°C, supernatant was removed, and the pellet was resuspended in 200-400 μL of cold 5% donkey serum (Jackson ImmunoResearch 017-000-12) in 0.1% PBS-Triton, transferred to a deep, V-bottom 96-well plate (Simport T11030), and incubated on an orbital shaker (80 rpm) for 1 hour at room temperature. After blocking, the 96-well plate was centrifuged at 70g for 5 minutes, supernatant was removed without disturbing the organoids, and 100 μL of primary antibodies diluted 1:100 in a solution of 5% donkey serum in 0.1% PBS-Triton was added to each well/stain group. [KC3] The organoids were incubated with primary antibodies (see Supplemental Table 2) for 24-72 hours at 4°C with mild rocking. Organoids were then centrifuged at 70g for 5 minutes, supernatant was removed, and organoids were washed with organoid washing buffer for 1-3 hours while orbitally shaking at 80 rpm. The washing procedure was repeated three times. Secondary antibodies (see Supplemental Table 2) diluted 1:200 in a solution of 5% donkey serum in 0.1% PBS-Triton were added and incubated overnight at 4°C protected from light with mild rocking. The following day, organoids were centrifuged at 70g for 5 minutes, supernatant was removed, and organoids were resuspended in DAPI diluted 1:2000 in 0.1% PBS-Triton, and incubated on an orbital shaker (80 rpm) for 15 minutes in the dark at room temperature. Organoids were then centrifuged at 70g for 5 minutes, supernatant was removed, and organoids were washed with organoid washing buffer for 1-3 hours while shaking at 80 rpm protected from light. The washing procedure was repeated three times before proceeding to the clearing procedure. Organoids were centrifuged at 70g for 5 minutes and the pellet was directly aspirated and transferred into a 1.5 mL tube. Thirty to 35 μL of fructose-glycerol solution (60% vol/vol glycerol [Sigma G7893] + 2.5 M fructose [Sigma F3510]) was added to the tube per desired mounted slide, gently resuspended, and allowed to optically clear overnight at 4°C protected from light. Before mounting, the organoids in fructose-glycerol solution were allowed to come up to room temperature to reduce viscosity during the mounting procedure. An ImmEdge Hydrophobic Barrier PAP pen (Vector Laboratories H-4000) was used to draw a 1 x 3.5-cm rectangle in the center of a microscope slide and a 1-cm-long piece of double-sided tape was placed at opposite ends of the rectangle. A cut 200 μL pipette tip was used to place 30-35 μL of organoids suspended in fructose-glycerol in the middle of the rectangle, and a 24 x 50 mm no. 1.5 coverslip (Globe Scientific 141515) was gently placed on top, resting on both pieces of double-sided tape. The mounted slides were stored flat at 4°C until ready to be imaged.

### Multi-photon confocal imaging of organoids

Wholemount immunostained organoids were imaged with a Nikon A1R inverted LUNV fitted with a 40X Plan Apochromat λ S Silicone Immersion objective (Nikon MRD73400). Using Nikon silicone immersion oil (refractive index of 1.41), a resonant Z-stack image of each organoid was acquired with live Nikon Ai Denoising and an averaging speed of 2-4. Images were processed and cropped in FIJI (FIJI/ImageJ v2.14) with minimal global adjustment of LUTs for acquired channels. For all resonant organoid images, an unsharp mask filter was applied in post with a radius (sigma) of 1 pixel and mask weight of 0.40-0.60 to improve image clarity.

### Mouse organoid plate imaging

For whole-well imaging of live organoids, plates were fed into a Cytation 10 Imager (BioTek, C10IPHC2) equipped with a CO_2_ gas controller (BioTek, 1210012). During imaging periods, plates were maintained at 5% CO_2_ and 37°C using *Cytation Gen5 Microplate Reader and Imager Software* (BioTek, version 3.08.01). Automated brightfield and fluorescent (GFP or RFP) 4x tile scans at 9 z-steps (∼50 um/step) were captured using plate-specific protocols (Falcon 24-well companion plate [Corning 353504] and Falcon Transwell Insert/Permeable Support with 0.4 um membrane [Corning 353095]). Stitched tile scans at 4x were used to generate z-projections for further quantification.

### Mouse organoid whole well quantification

Stitched 4x Z-projections from each well were uploaded into a custom FIJI macro (executed in FIJI/ImageJ v2.14) to quantify the number of organoids per well using brightfield, GFP, or RFP filters. The macro facilitated batch analysis for each experiment, minimizing the subjectivity of hand counting. In brief, given specific input parameters, the macro contained commands to: set the scale based on the diameter of each transwell, perform convoluted background subtraction, manually adjust image threshold, convert to mask, analyze and count particles meeting a specific threshold, and export table data. Data was imported into GraphPad Prism 10.0 for analysis. T-tests were used for comparison of 2 groups, and ANOVA with prespecified multiple comparisons was used to compare 3 or more groups.

### Human iAT2 Single Cell RNA sequencing (scRNA-seq)

For iAT2 cell profiling, 5 time points over a single 13-day passage cycle and an additional 6th time point 1 day after the subsequent passage were chosen based on proliferation kinetics to perform scRNAseq on the inDrops platform. At each indicated day post-passage, iAT2 cells (SPC2-ST-B2 clone) were dissociated with 2mg/mL dispase II, followed by 0.05% trypsin-EDTA, and resuspended in wash buffer with Calcein Blue. The cells were sorted on live single cells, and 4,000 cells per time point were captured for inDrops (Harvard University Single Cell Core). Reads were demultiplexed using the protocol at https://github.com/indrops/indrops. Reads were then aligned using Bowtie/RSEM to a custom GRCh38 reference augmented with the tdTomato reporter. Gene-by-cell UMI count matrices were compiled per sample and imported into Seurat for individual processing, merging and further analysis according to recently published methods^30^ as indicated in the text with dimensionality reduction visualization performed using SPRING^52^.

### Mouse Lung scRNA-seq, Alignment & Quality Control

For *Sftpc*^creERT2^ R26R^DTA^ mouse cells, mouse lungs were digested as previously described^53^. Briefly, lungs were inflated with 1.5 mL of digestion buffer (9.5U/mL Elastase, 20U/mL Collagenase, 5U/mL Dispase), incubated at 37C, and passed through 70µm and 40µm cell strainers. Cells were stained with and sorted on EpCAM, CD31, and CD45 antibodies, along with DRAQ7 viability dye. Cells were captured for 10X 3’v3 processing with targeted cell recovery of 10,000 combined cells at the Boston University Single Cell Sequencing Core.

From Eed^CKO^ mouse single cell suspension, the maximum number of cells were loaded into a single channel of the Chromium system using the 3v4 single cell reagent kit (10X Genomics, Pleasanton, CA) by the Cincinnati Children’s Hospital Medical Center Single-Cell Gene Expression Core.

Raw sequencing data (submitted to Gene Expression Omnibus) were aligned to the Mus musculus reference mm10 with Cell Ranger 4.0.3 (10X Genomics), generating expression count matrix files. Cells with fewer than 500 features or greater than 5000 features, as well as cells that contained greater than 25% of reads from mitochondrial genes, were removed. Putative multiplets were removed using DoubletFinder^54^ (version 2.0).

### Mouse Lung scRNA-seq Data Analysis

The Seurat package (version 5, https://satijalab.org/seurat/)^55^ in R 4.4.3 was used for identification of common cell types across different experimental conditions, differential expression analysis, and most visualizations. Libraries from different samples were integrated with Harmony^56^. Integrated data was scaled, and regression performed for mitochondrial genes, ribosomal genes, and cell cycle state. Commands used included NormalizeData, FindVariableFeatures, ScaleData, RunPCA, RunUMAP, FindNeightbors, and FindClusters. Manual annotation of cellular identity was performed by via identification of differentially expressed genes for each cluster using Seurat’s implementation of the Wilcoxon rank-sum test (FindMarkers()) and comparing those markers to known cell type-specific genes from LungMAP^57^. These Seurat objects were used as the basis for additional analytical tools to explore our scRNAseq data. We generated pseudotime trajectories and gene regulation across pseudotime with Monocle3 (https://cole-trapnell-lab.github.io/monocle3/)^58^. Visualizations were generated by these tools, with additional use of ggplot2 when needed.

### Statistical Tests

All data met the assumptions of the statistical tests used. Statistical tests used for single-cell analyses are described in relevant section. For comparing the differences between groups, we used either unpaired two-tailed Student’s *t*-test for two groups or ANOVA with prespecified multiple comparison testing for groups of three or more (. These statistical analyses were performed in GraphPad Prism 10.0. Details of statistical tests are reported in each figure legend. A *P* value of less than 0.05 was considered significant; \**P* < 0.05, \*\**P* < 0.01, \*\*\**P* < 0.001, and \*\*\*\**P* < 0.0001

## TABLES

**Supplemental Table 1:**
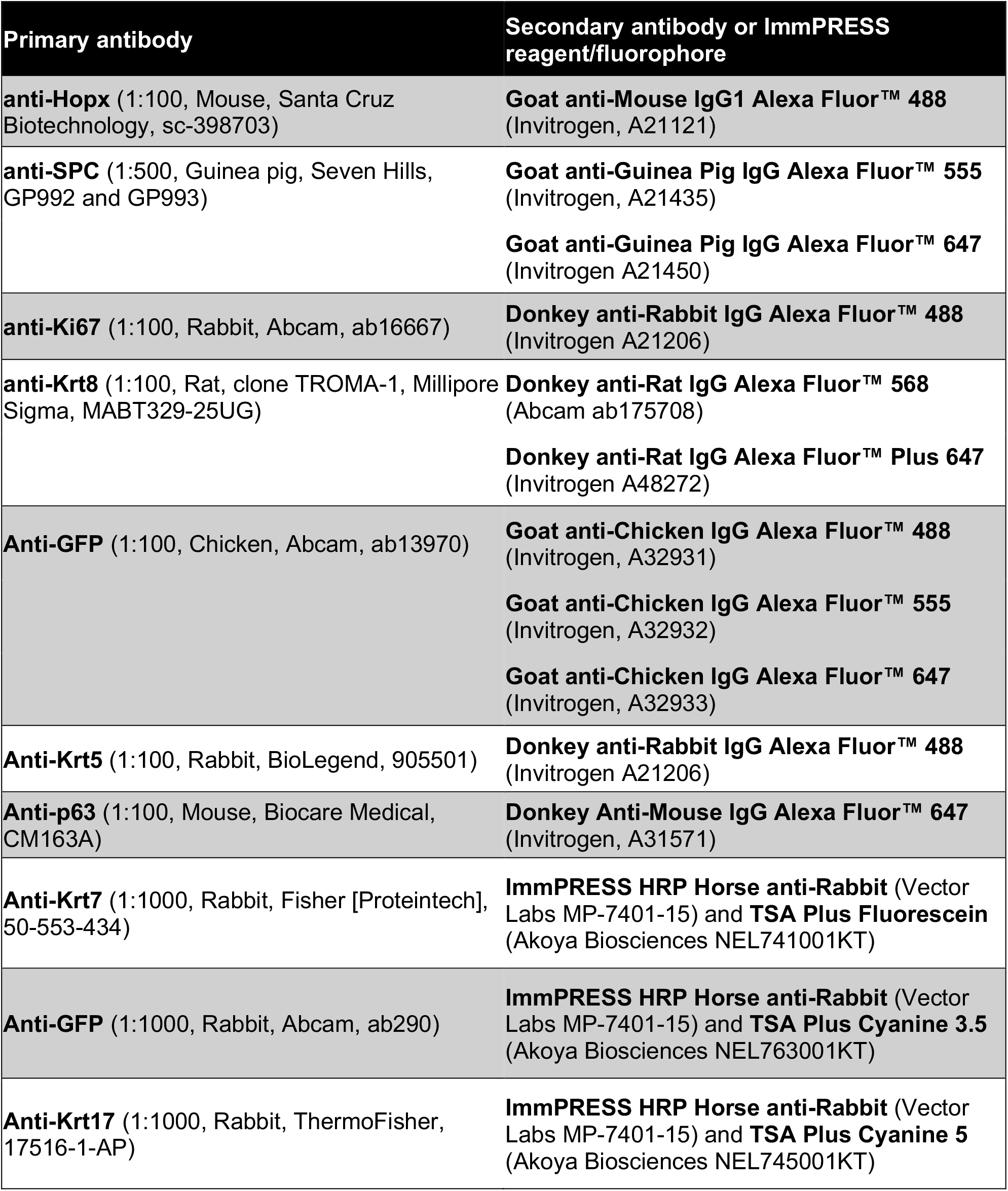
Antibodies used for immunostaining on lung sections.

**Supplemental Table 2:** Antibodies used for wholemount organoid immunostaining.

| **Primary antibody (1:100)** | **Secondary antibody (1:200)** |
| --- | --- |
| **anti-RAGE** (Rat, R&D Systems, MAB1179) | **Donkey anti-Rat IgG Alexa Fluor™ 488**<br>(Invitrogen, A21208)<br>**Donkey anti-Rat IgG Alexa Fluor™ Plus 647** (Invitrogen A48272) |
| **anti-SPC** (Guinea pig, Seven Hills, GP992 and GP993) | **Goat anti-Guinea Pig IgG Alexa Fluor™ 555** (Invitrogen, A21435)<br>**Goat anti-Guinea Pig IgG Alexa Fluor™ 647** (Invitrogen A21450) |
| **anti-Ki67** (Rabbit, Abcam, ab16667) | **Donkey anti-Rabbit IgG Alexa Fluor™ 488**<br>(Invitrogen A21206)<br>**Donkey anti-Rabbit IgG Alexa Fluor™ 647**<br>(Invitrogen A31573) |
| **anti-Krt8** (Rat, clone TROMA-1, Millipore Sigma, MABT329-25UG) | **Donkey anti-Rat IgG Alexa Fluor™ 488**<br>(Invitrogen A21208)<br>**Donkey anti-Rat IgG Alexa Fluor™ Plus 647** (Invitrogen A48272) |
| **anti-Krt5** (Rabbit, BioLegend, 905501) | **Donkey anti-Rat IgG Alexa Fluor™ 488**<br>(Invitrogen A21208) |

## SUPPLEMENTAL MATERIALS

**Extended Data Figure 1.**
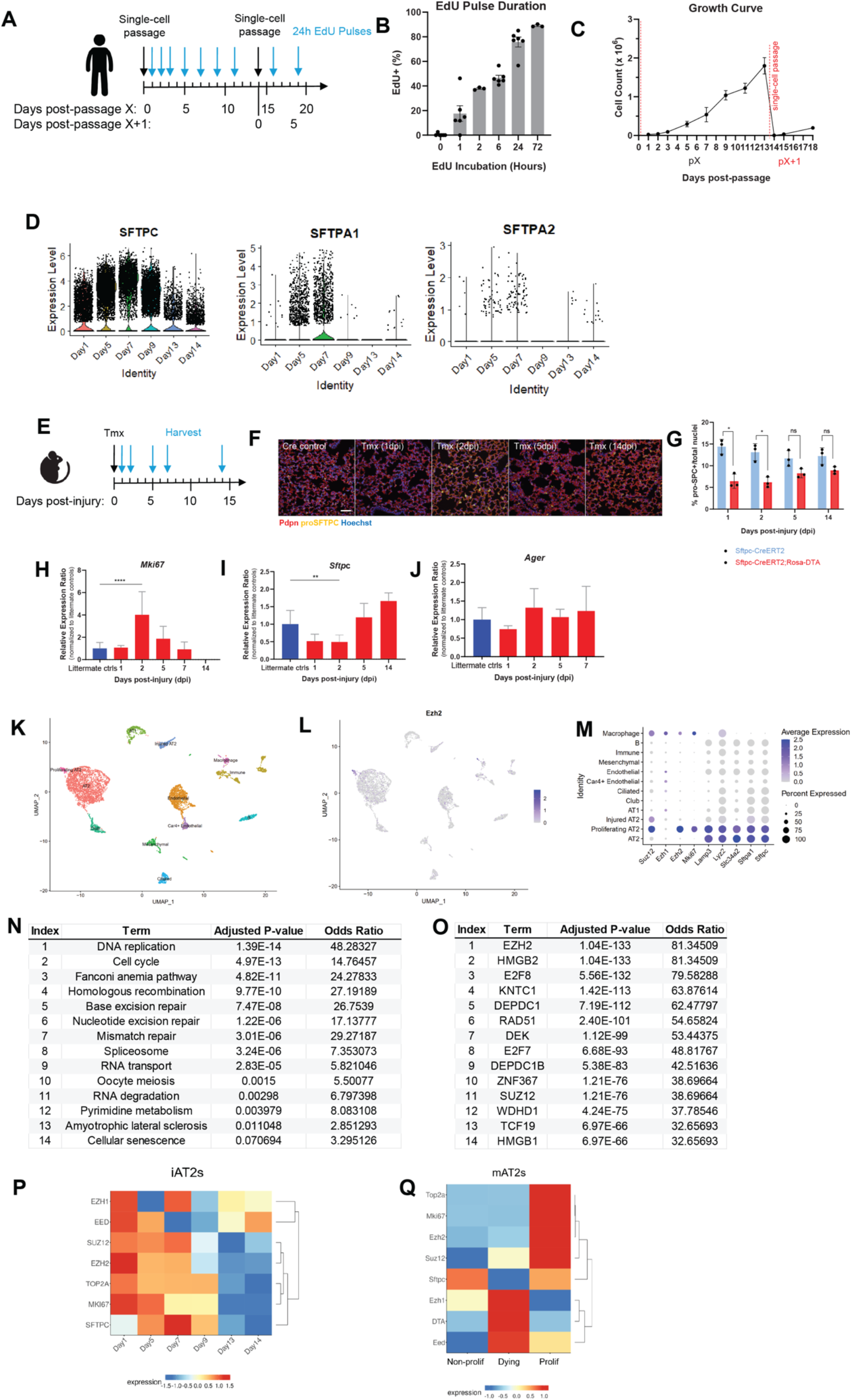
Single cell analysis of proliferating AT2 cells in mouse and human. (A) Experimental design for human iAT2 timepoints used for single cell sequencing. (B). Percentage of proliferating AT2 cells by Edu labeling. (C) Growth curve for total cells in iAT2 culture at each day post passage. Passage is denoted by dotted line, and next passage is denoted in red. (D)Violin plots showing the expression of the AT2 cell maturation marker genes *SFTPC, SFTPA1*, and *SFTPA2* at each timepoint. (E) Experimental design for mouse diphtheria-toxin (DT) mediated ablation of AT2 cells. (F) IHC showing AT1 cells marked by Pdpn and AT2 cells marked by pro-SPC at each timepoint following induction of DT ablation. (G) Percentage of pro-SPC positive cells normalized to total nuclei at each timepoint following induction of DT. (H) QPRC showing expression of the proliferation marker *Mki67*, the AT2 marker *Sftpc*, and the AT1 marker *Ager* at each timepoint following DT-mediation AT2 ablation. (K-M) Single cell RNA sequencing results from DT-mediated AT2 ablation, with UMAP showing cell type clustering (K), UMAP showing distribution of expression of *Ezh2* mRNA (L), and dot plot showing PRC2 complex and AT2 marker mRNA expression per cell type (M). (N-O) KEGG pathway enrichment analysis (N) and upstream regulatory prediction (O) for genes enriched in proliferating AT2 cells (as identified in Figure 1H). (P-Q). Heatmaps showing PRC2 and proliferative gene expression per day post-iAT2 passage (P) or based on cell cycle state in murine AT2 after DTA-mediated AT2 ablation (Q).

**Extended Data Figure 2.**
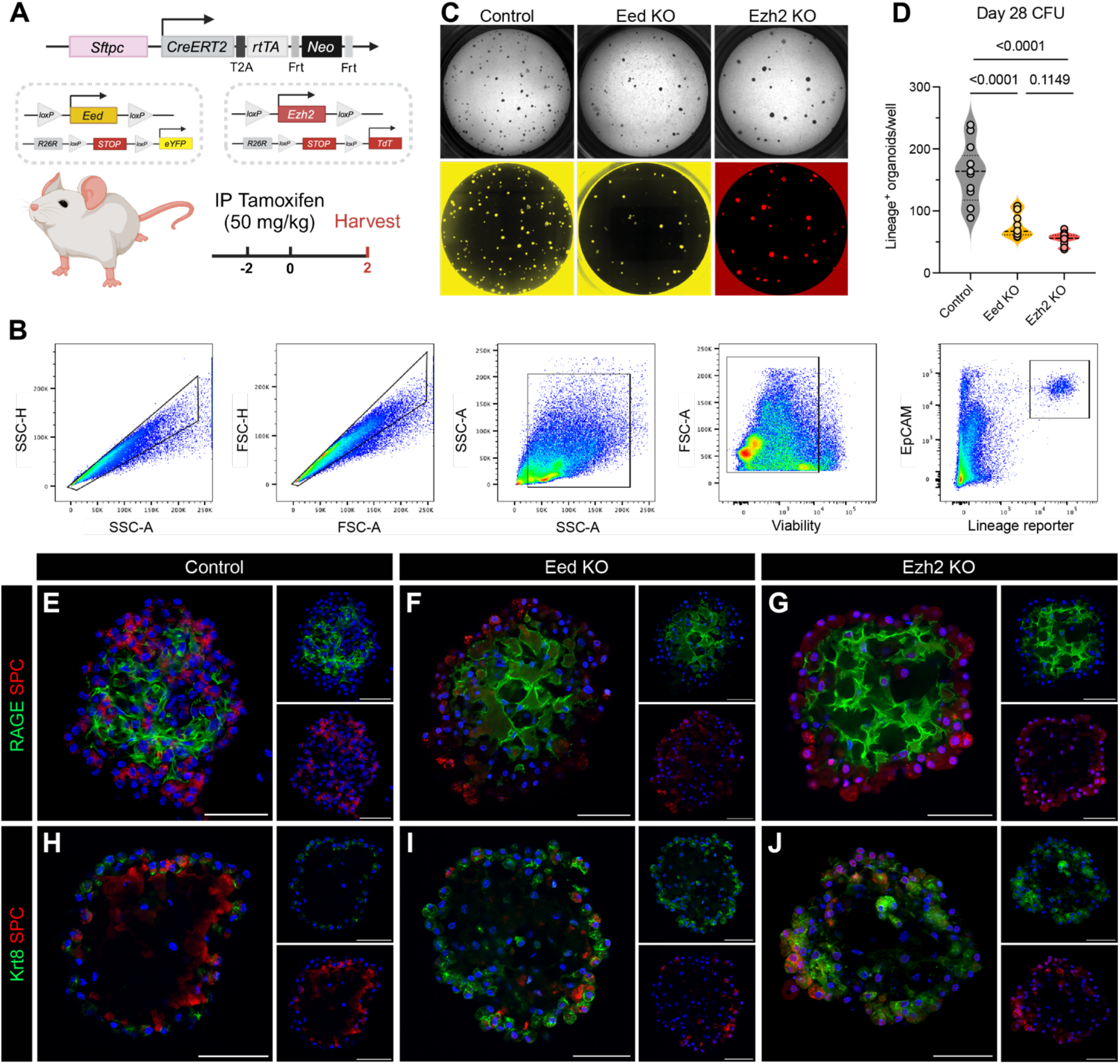
Impact of PRC2 complex loss of function in murine alveolar organoids. (A) Schematic detailing the generation of a SPC-driven tamoxifen-inducible Eed or Ezh2 floxed mouse with fluorescent lineage reporter, followed by tamoxifen treatment regimen used for organoid sort. (B) Gating strategy used to sort AT2 cells following *in vivo* induction of tamoxifen. Cells sorted in this manner were used to initiate organoid culture. (C) Brightfield (top) and fluorescent (bottom) well images of control, Eed KO, and Ezh2 KO organoids after 28 days of culture. (D) Quantification of organoid colonies measured from fluorescent well images. Dots represent technical replicates from n = 2 mice per group, dashed lines represent medians, and dotted lines represent quartiles of individual datasets. **** = p < 0.0001 by ANOVA with Multiple Comparisons. (E-J) Wholemount IF images of representative organoids from control (E,H), Eed^CKO^ (F,I), and Ezh2^CKO^ (G,J) cultures stained for RAGE and SPC (top row) or Krt8 and SPC (bottom row). For E-J, scale bars = 50 µm.

**Extended Data Figure 3.**
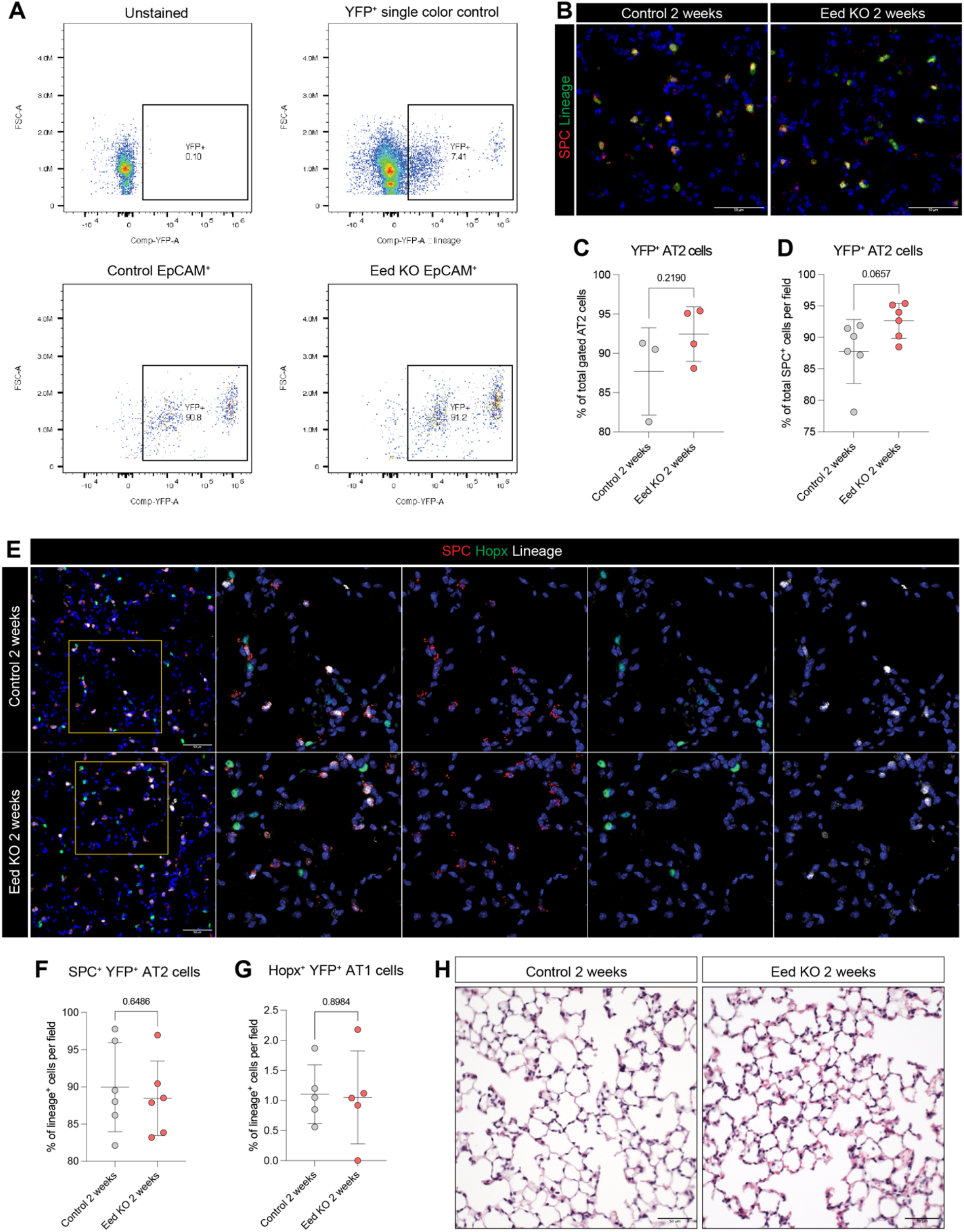
AT2 population dynamics two weeks post-Eed KO. (A) Identification of YFP^+^ cells by flow cytometry 2 weeks post-tamoxifen treatment. Gates show YFP lineage reporter expression relative to cell size (FSC). (B) Identification of YFP^+^ AT2 cells (YFP^+^ SPC^+^) by IHC 2 weeks post-tamoxifen treatment. (C) Quantification of YFP^+^ cells described in A. Dots represent individual biological replicates from n = 3-4 mice per group. Bars represent means +/-SD. Statistical analysis by unpaired t-test. (D) Quantification of YFP+ AT2 cells described in B. Dots represent individual biological replicates from n = 6 mice per group. Bars represent means +/-SEM. Statistical analysis by unpaired t-test. (E) Representative IF images from mice described above stained for SPC, Hopx, and YFP lineage reporter. (F) Quantification of SPC^+^ YFP^+^ AT2 cells as a proportion of total YFP^+^ cells from mice in E. Dots represent individual biological replicates from n = 6 mice per group. Bars represent means +/-SEM. Statistical analysis by unpaired t-test. (G) Quantification of Hopx^+^ YFP^+^ AT1 cells as a proportion of total YFP^+^ cells from mice in E. Dots represent individual biological replicates from n = 5 mice per group. Bars represent means +/-SEM. Statistical analysis by unpaired t-test. (H) Representative H&E images of control and Eed KO mice described above.

**Extended Data Figure 4.**
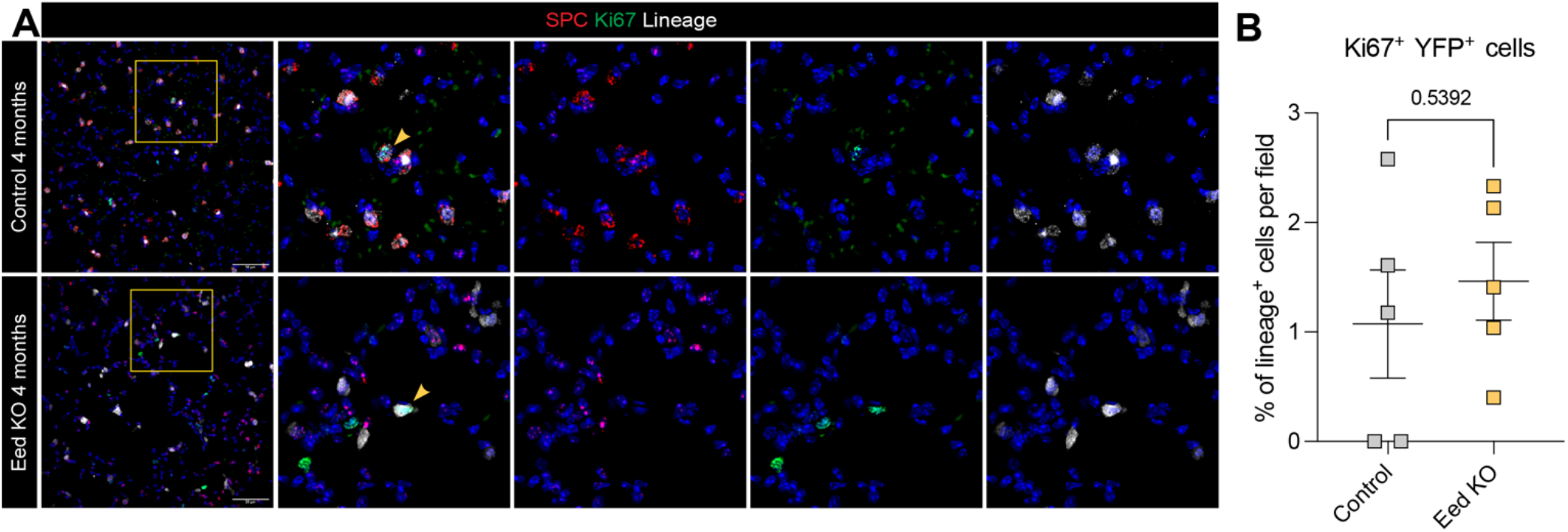
Evaluation of AT2 proliferation dynamics following EED loss of function. (A) Representative IF images from mice 4 months post-tamoxifen treatment stained for SPC, Ki67, and YFP lineage reporter. (B) Quantification of cells described in A. Dots represent individual biological replicates from n = 5 mice per group. Bars represent means +/-SEM. Statistical analysis by unpaired t-test.

**Extended Data Figure 5.**
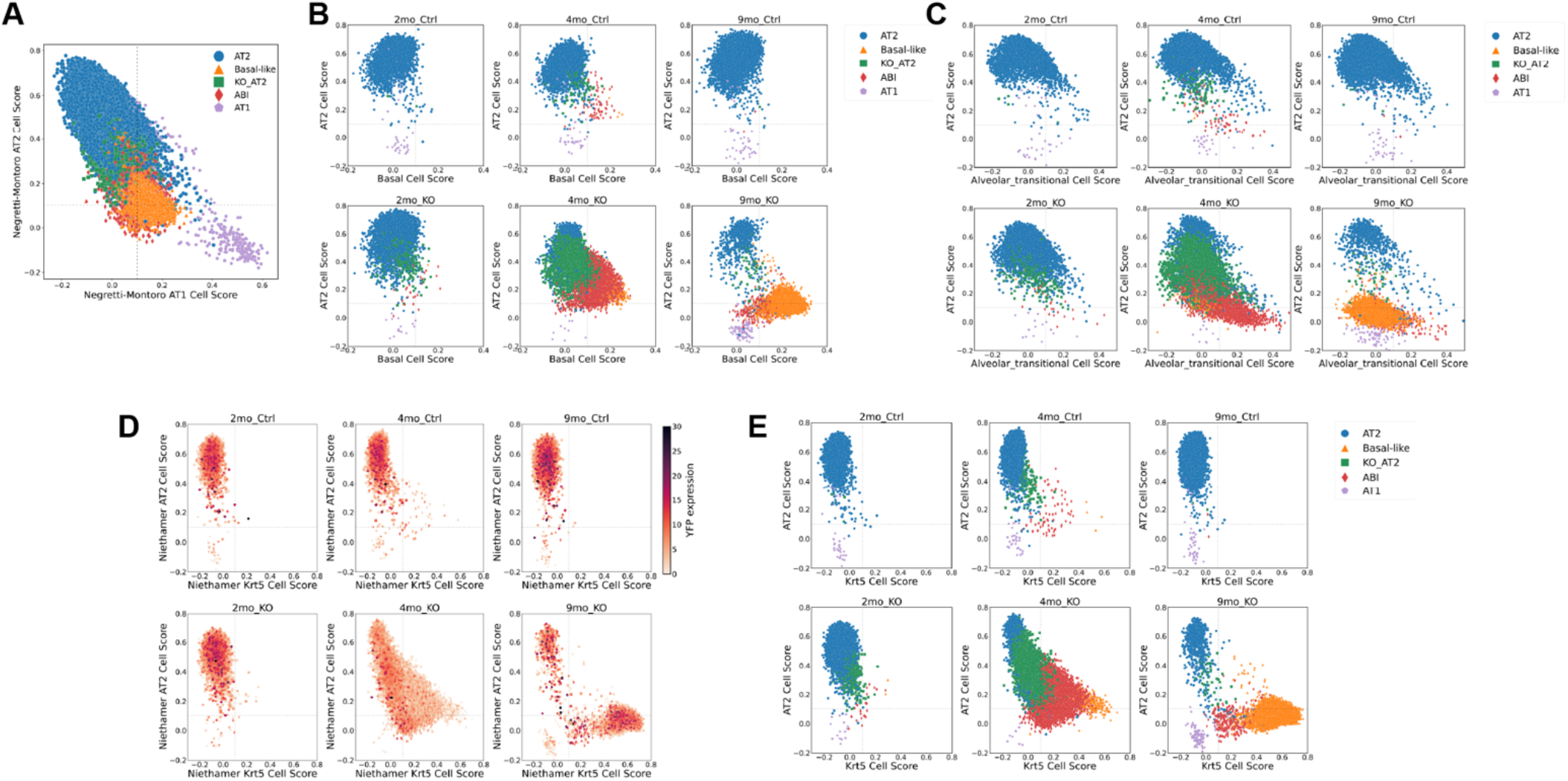
scTOP analysis of cell state transition following EED loss of function *in vivo*. (Similarity score of cell states identified in murine scRNAseq (see Figure 4) compared with cell identities from uninjured cells from Negretti et al^38^ and Montoro et al^39^ scRNAseq atlases. (A) Cells from all timepoints plotted on axis comparing AT2 cells score and AT1 cells score from uninjured lung atlases. (B) Cells from each timepoint and condition plotted on axis comparing AT2 cell score and basal cell score in the uninjured lung atlases; compare to Figure 4M. (D) Cells from each timepoint and condition plotted on axis comparing AT2 cell score and Alveolar_transitional cell score in the post-influenza lung injury atlas^40^; compare to Figure 4N. (D-E) Cells from each timepoint and condition plotted on axis comparing AT2 cell score and Krt5 cell score in the injury atlas; compare to Figure 4O. Cells in D are colored based on EYFP expression, confirming that cells in the EED^CKO^ lineage continue to express the lineage marker despite transitioning to basal-cell like state.

**Extended Data Figure 6.**
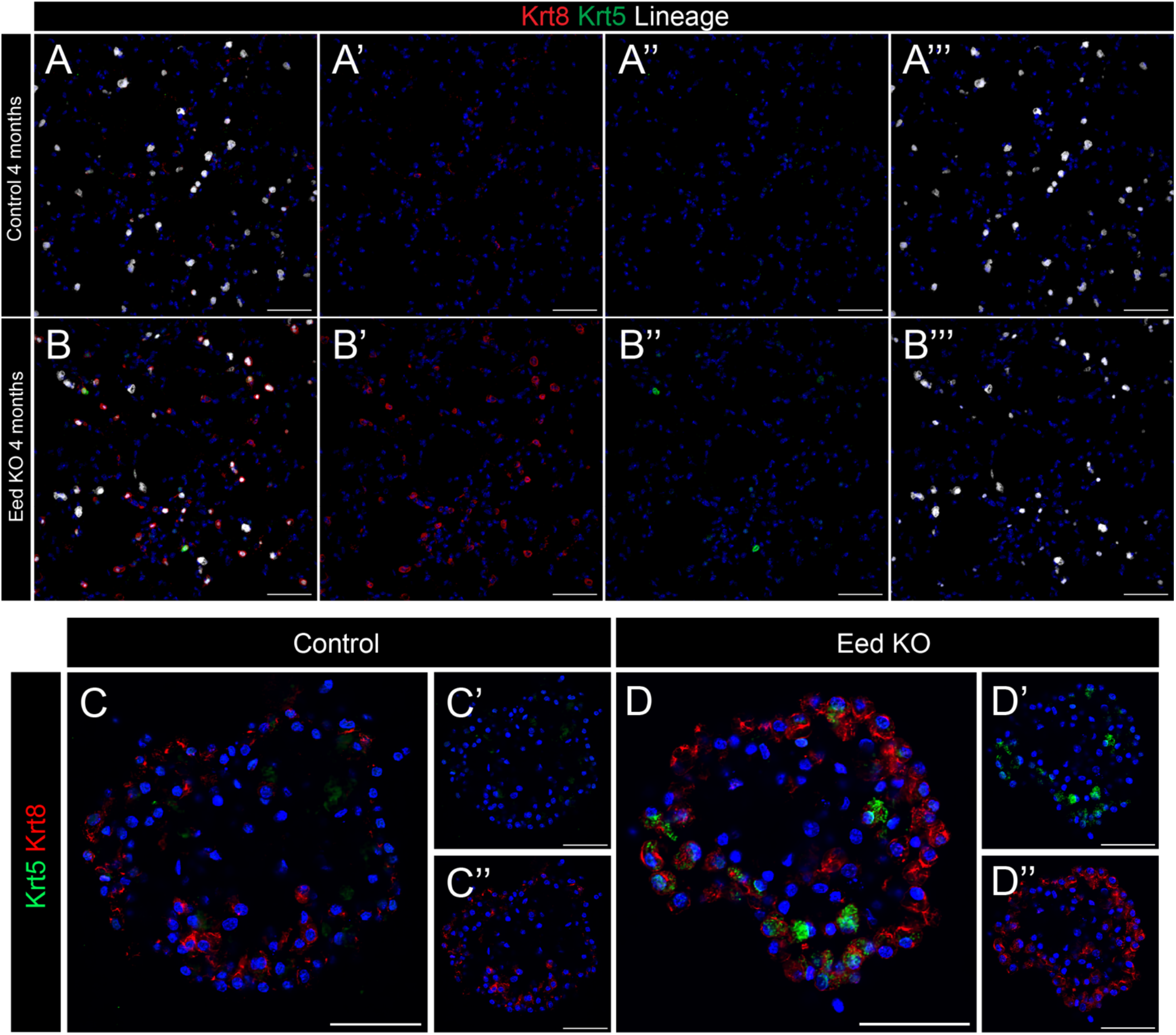
Time dependent acquisition of Krt5 expression in EED^CKO^ lineage. (A-B) Expression of Krt8 and Krt5 in lineage labeled AT2 cells at 4 months post tamoxifen demonstrating widespread acquisition of Krt8^high^ cell state and rare Krt5^pos^ cell state in the EED^CKO^ lineage. (C-D) Expression of Krt8 and Krt5 in alveolar organoids derived from EED^CKO^ AT2 cells at 28d of culture showing more rapid acquisition of Krt8^high^ and Krt5^pos^ cell states in the proliferative expansion of EED^CKO^ lineage cells in lung organoid formation. For A-B, scale bars = 200 µm; for C-D, scale bars = 50 µm.

